# The resilience of *Salmonella* to bile stress is impaired due to the reduced efflux pump activity mediated by the antioxidant enzyme YqhD

**DOI:** 10.1101/2024.11.23.625033

**Authors:** Kirti Parmar, Yogyta Kumari, Raju S Rajmani, Dipshikha Chakravortty

**Affiliations:** Department of Microbiology and Cell Biology, Division of Biological Sciences, Indian Institute of Science, Bangalore, India; Centre of Infectious Disease Research, Indian Institute of Science, Bangalore, India; Adjunct Faculty, School of Biology, Indian Institute of Science Education and Research, Thiruvananthapuram, India

**Author notes:** Corresponding author: Prof. Dipshikha Chakravortty.

## Abstract

Bile salts play a critical role in modulating the host gut. They have antimicrobial properties wherein they disrupt the bacterial membrane and produce reactive oxygen species, causing DNA damage. Bile-resistant pathogen like *Salmonella* regulates their metabolic activity to counteract the effects of bile. This study explores the role of YqhD, a putative alcohol dehydrogenase, in *Salmonella’*s bile salt susceptibility. We observed increased survival of *yqhD* mutant in the in vitro studies in LB media with bile, liver cell line HepG2 and C57BL/6 mice on treatment with 8% sodium cholate in the intestine. Bile salts are produced for the digestion of fat. Replacing the chow diet with a high-fat diet (HFD) in mice increased organ burden in C57BL/6 mice of the *yqhD* mutant.

Mutation of *yqhD* on bile salt exposure leads to increased reactive oxygen species and modulation of antioxidant genes in the bacteria. The oxidative stress of the *yqhD* mutant is indispensable for improved survival when exposed to bile salt. The addition of antioxidant glutathione *in vitro* reduced the enhanced growth of the *yqhD* mutant. Similarly, in the gp91^−/−^ phox mice, there was a reduced organ burden of *yqhD* mutant on exposure to HFD compared to chow-fed mice. Furthermore, the *yqhD* mutant increases AcrAB efflux pump activity regulated by RamA/R regulon.

**Teaser:** Redox gene *yqhD* regulates bile salt susceptibility by modulating *Salmonella’s* ROS and efflux pump activity.

**Graphical abstract:** **A.** STM Δ*yqhD* has less organ burden in the C57BL/6 mice than STM WT, and there was a similar organ burden in *gp*91^phox−/−^. On HFD exposure, the organ burden of STM Δ*yqhD* became similar to STM WT in C57BL/6 mice, and there was reduced organ of STM Δ*yqhD* burden when *gp*91^phox−/−^ were exposed to HFD.
**B.** On bile salt exposure, YqhD in *Salmonella* reduces ROS in bacteria and expression of RamA, decreasing the AcrAB efflux pump activity and reducing survival.

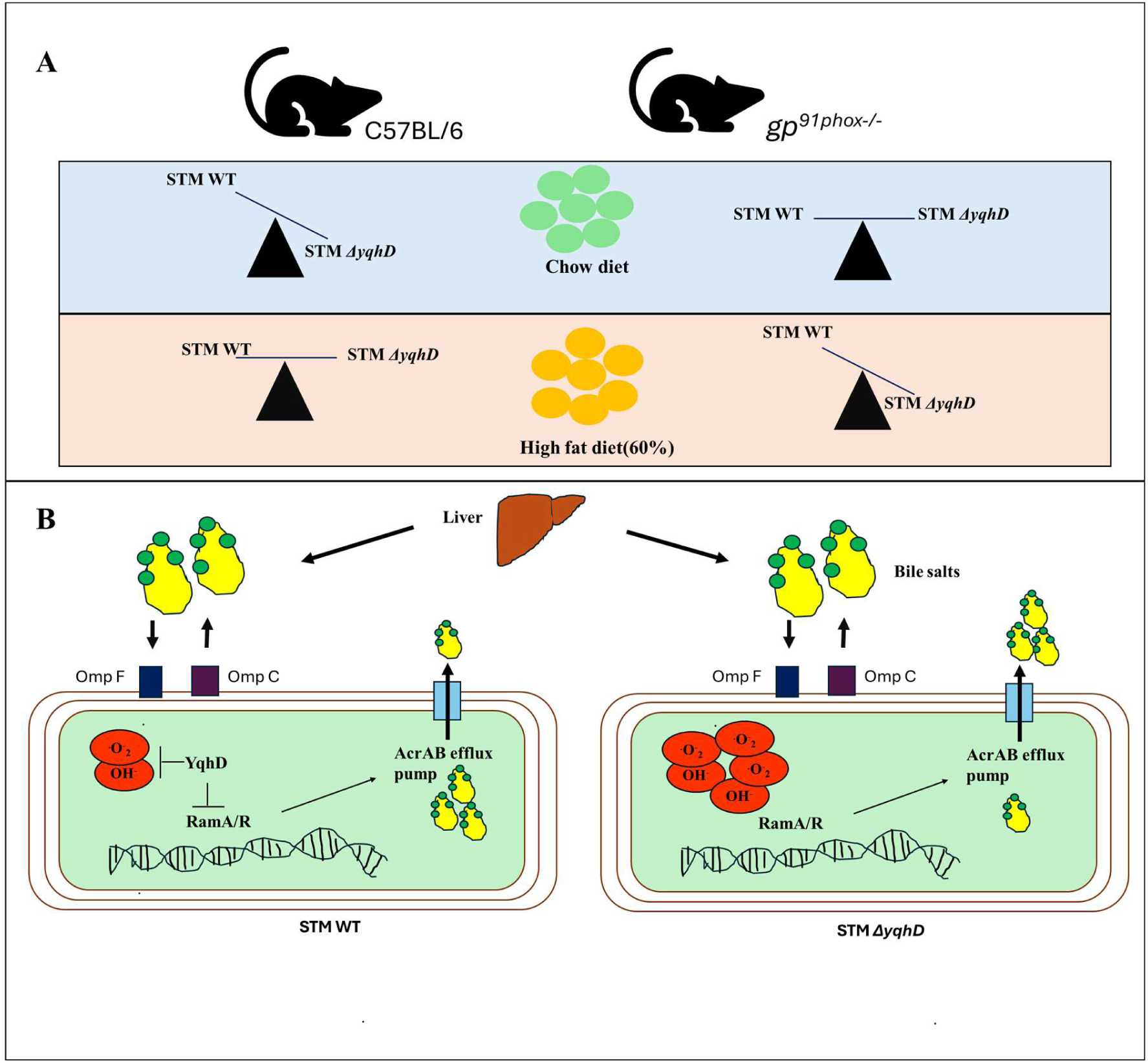

## Introduction

*Salmonella* is a Gram-negative, rod-shaped bacterium from the *Enterobacteriaceae* family. It infects a variety of hosts. *Salmonella* Typhimurium is the second most common serovar isolated from humans and has been used in this study(*1*). *Salmonella* is a foodborne pathogen facing multiple detrimental host factors and environments while passing through the host’s gastrointestinal tract(*2*). One of the biggest challenges *Salmonella* faces is bile salts.

Bile is a digestive secretion synthesised by hepatocytes in the liver from cholesterol and stored in the gall bladder or directly secreted into the intestine. Bile salts are a major component of bile and are potent detergents. Physiologically, bile aids in the digestion of fats and the absorption of fat-soluble vitamins(*3*). Although antibacterial, they are used as environmental signals by bacteria to regulate their pathogenesis and bile resistance(*4–8*). *Salmonella enterica* is an example of a bile-resistant pathogen; it colonises and persists in the gall bladder during chronic infection(*9*). Mary Mallon was the first known case of chronic infection; she was an asymptomatic carrier and transmitted bacteria in every household where she worked as a cook(*10*). Although enteric bacteria have been known to have bile resistance, new genes and the molecular mechanism responsible for bile resistance are still elusive and require further investigation.

In our study, we investigated the role of YqhD in *Salmonella enterica* serovars Typhimurium and Typhi. YqhD is an NADPH-dependent aldehyde reductase with zinc as a cofactor(*11*). Studies have shown that YqhD decreases Reactive oxygen species(ROS) levels and ROS-mediated effects by acting on aldehydes such as propanal and butanal, which are generated upon membrane peroxidation and glyoxal that is produced by glucose oxidation (*12, 13*). Transcriptomics analysis study of *Salmonella* under varying stresses showed that *yqhD* was induced by iron deficiency, bile and osmotic stress(*14*). For the first time, we decipher the role of YqhD in potentiating bile susceptibility in *Salmonella*. We show that *yqhD* deletion leads to increased survival in the presence of bile, although it has greater ROS. Deleting *yqhD* leads to the induction of the AcrAB efflux pump through the RamA/R regulon, independent of the SoxR/S regulon. We delineate a novel mechanism determining sensitivity to bile in *Salmonella,* involving YqhD as a critical player by modulating the efflux pump activity through RamA.

## Results

### 1. *yqhD* gene reduced *Salmonella* survival on exposure to bile salts

To study the function of YqhD in *Salmonella,* we observed the *yqhD* mRNA expression in STM WT at various time points. We observed that expression was highest at 3 hours and reduced significantly with time in LB media (supplementary Fig.1A). On treatment with 7%bile salts in LB media to STM WT, *yqhD* expression was decreased significantly (Fig 1A).

**FIG 1.**
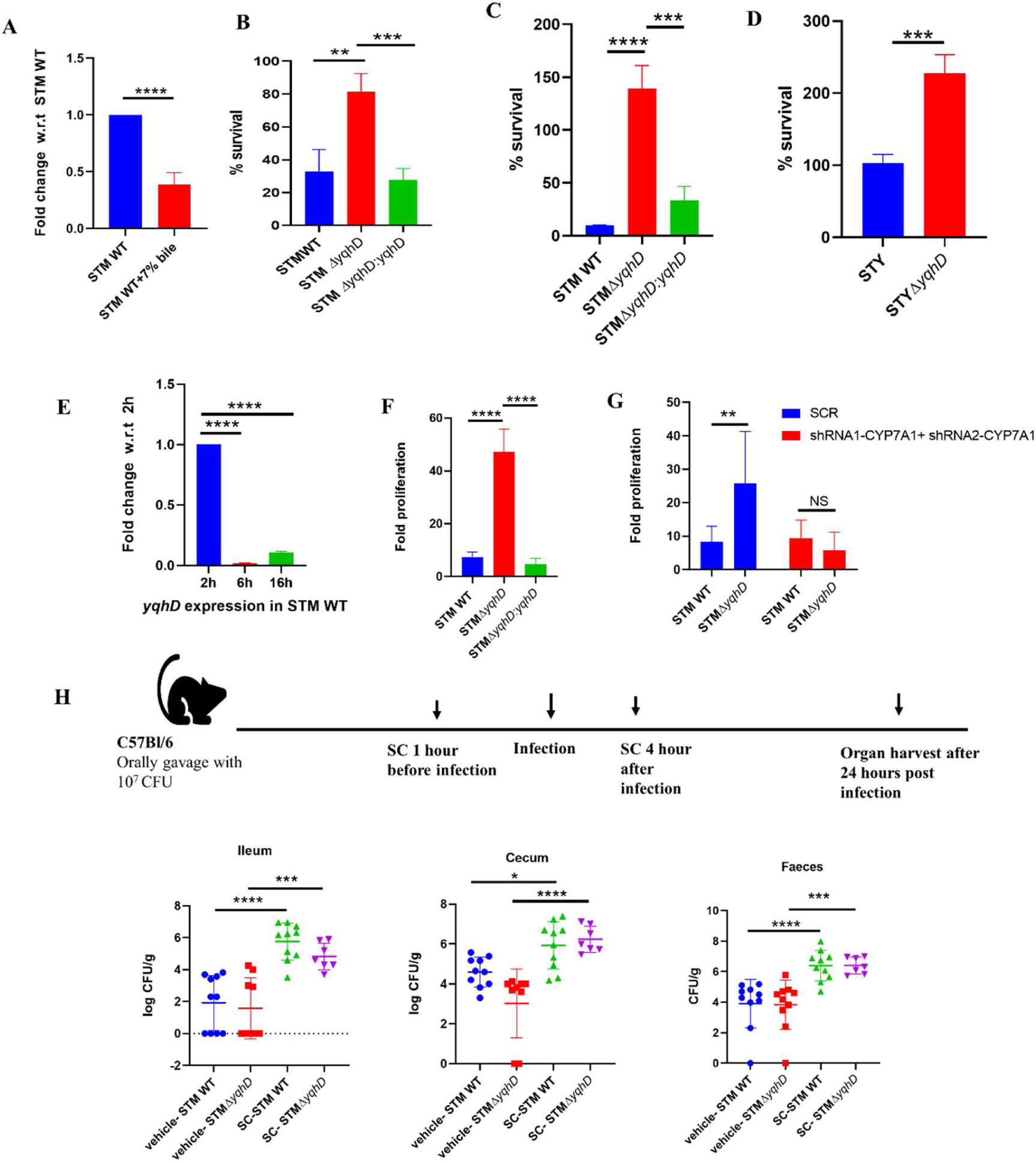
Deletion of *yqhD* in *Salmonella* increases growth on exposure to bile salts. **A**. The mRNA expression of *yqhD* in STM WT on treatment with 7% bile with respect to untreated. **B**. The *in vitro* sensitivity assay on the treatment of 7% primary bile salt (Himedia) to STM WT, STMΔ*yqhD* and STMΔ*yqhD:yqhD* normalised to un-treated by CFU/ml (N=5,n=2). **C**. The *in vitro* sensitivity assay on treatment on the treatment of 1 % sodium deoxycholate (Sigma) to STM WT, STMΔ*yqhD* and STMΔ*yqhD:yqhD* by CFU/ml (N=3,n=2). **D**. The *in vitro* sensitivity assay on the treatment of 7% primary bile salt (Himedia) to *Salmonella* Typhi CT18, *Salmonella* Typhi Δ*yqhD* by CFU/ml (N=3,n=2). **E**. The mRNA expression of *yqhD* in STM WT upon infection in the HepG2 cells with respect to STM WT. **F**. Fold proliferation of STM WT, STMΔ*yqhD* and STMΔ*yqhD:yqhD* in HepG2 cells (N=5,n=3). Analysis by unpaired two-tailed Student’s t-test; p values****<0.0001, ***<0.001, **<0.01, *<0.05. **G**. Fold proliferation STM WT, STMΔ*yqhD* and STMΔ*yqhD:yqhD* in the CYP7A1 knockdown in HepG2 cells. Analysis performed using two-way ANOVA; p values****<0.0001, ***<0.001, **<0.01, *<0.05. **H**. Representation of animal experiment methodology in C57Bl/6 male mice, ten animals per cohort were taken, and bacterial organ burden in ileum, cecum and faeces in mice was determined. Mann-Whitney test was performed for statistical analysis.; p values****<0.0001, ***<0.001, **<0.01, *<0.05. All data are represented as mean±SD. Data is from one experiment representative of N ≥3 independent experiments.

To further study the role of *yqhD* in *Salmonella* Typhi and Typhimurium, we generated a *yqhD* deleted strain using the lambda red recombinase system (*15*). We found that the deletion of *yqhD* did not alter its *in vitro* growth in LB media (supplementary fig.1B, C) and when examining the effect of various stresses reported to induce *yqhD.* We did not observe a change in the growth of STM *ΔyqhD* in the presence of iron chelator 2,2’-bipyridyl and osmotic stress inducer sodium chloride compared to STM WT. While STM *ΔyqhD* exhibited a higher bacterial growth in the presence of the bile salt mixture (supplementary Fig.1 D-F). We were further intrigued to look into the role of *yqhD,* an antioxidant gene, in regulating the bile salt response in *Salmonella*.

The minimal inhibitory concentration(MIC) of bile salts for *Salmonella* Typhimurium has been reported in the literature to be 7% for sodium deoxycholate and 14% for oxbile, respectively (*8*). We used a 7% bile salt mixture for most of our experiments. We observed an increase in the survival of STM *ΔyqhD* upon exposure to both primary (7%bile salts mixture) and secondary bile salt (1%sodium deoxycholate) on analysis of CFU/ml (Fig 1B, C). Similarly, for *Salmonella* Typhi, there was increased survival on exposure to 7% bile for STY *ΔyqhD* (Fig 1D).

Bile is secreted by hepatocytes in the liver. Thus, we validated our observation in the hepatocyte cell line HepG2. We observed that upon infection of STM WT, the mRNA expression of *yqhD* was reduced over time (Fig 1E). Upon performing a gentamicin protection assay, we observed that STM *ΔyqhD* shows a hyperproliferation compared to STM WT (Fig1F). Bile is produced from cholesterol in hepatocytes, with CYP7A1 (cytochrome P450 family 7 subfamily A member 1) catalysing a rate-limiting step (*16*). To confirm our findings that bile is responsible for the increased proliferation of STM Δ*yqhD* in the HepG2 cells. We performed lentivirus-mediated knockdown of CYP7A1 in HepG2 cells. We confirmed it using q-RT PCR and obtained 50% knockdown (Supplementary Fig 2A). We further confirmed it by measuring bile in the media and HepG2 cells. There was a slight reduction in the bile concentration in both the media and cells (Supplementary Fig 2B). Upon infection of knockdown cells, we observed that the proliferation of STM Δ*yqhD* was similar to STM WT (Fig 1G). Thus, loss of *yqhD* generates resistance to bile salts and enhances the survival of *Salmonella* in the presence of bile salts and hepatocytes. Next, we examined the effect of sodium cholate, one of the major primary bile salts secreted by hepatocytes in the liver (*17*). 8% sodium cholate treatment in mice has been reported to increase colonisation of STM WT (*19*). Upon treatment with sodium cholate, we observed increased colonisation of STM Δ*yqhD* in the cecum by 2 log fold compared to 1 log fold increase in STM WT and reduced excretion of STM Δ*yqhD* (2.2 log fold) compared to STM WT (2.8 fold)( Fig. 1H). However, STM WT increased by 4 log fold in the ileum and STM Δ*yqhD* 2 log fold. These results show that exposure to sodium cholate in mice leads to increased organ burden of STM Δ*yqhD* in the cecum and concomitantly reduced excretion in faeces.

Furthermore, bile is a detergent, and it helps in the emulsification of lipids(*18*). Bile secretion is induced upon a high-fat diet for the absorption of fats. Studies show that a high-fat diet and bile salt secretion boost *Salmonella* gut colonisation (*18*). To examine the effect of a high-fat diet on colonisation of STM *ΔyqhD* in C57BL/6 mice., mice were kept on a high-fat diet (60%) or chow diet for 14 days, and their weight and food intake were measured (Supplementary Fig 3A). On the 15^th^ day, the mice were infected, and we observed that three days post-infection, the mRNA levels of *yqhD* were reduced in STM WT in mice on HFD compared to the chow diet (Fig 2B). Upon infection of STM *ΔyqhD,* there was no difference in organ burden in the primary sites of infection on the exposure to HFD. Colonisation of STM *ΔyqhD* was increased in secondary sites of infection liver and spleen when mice were kept on HFD compared to the chow diet (Fig 2A).

**Fig. 2.**
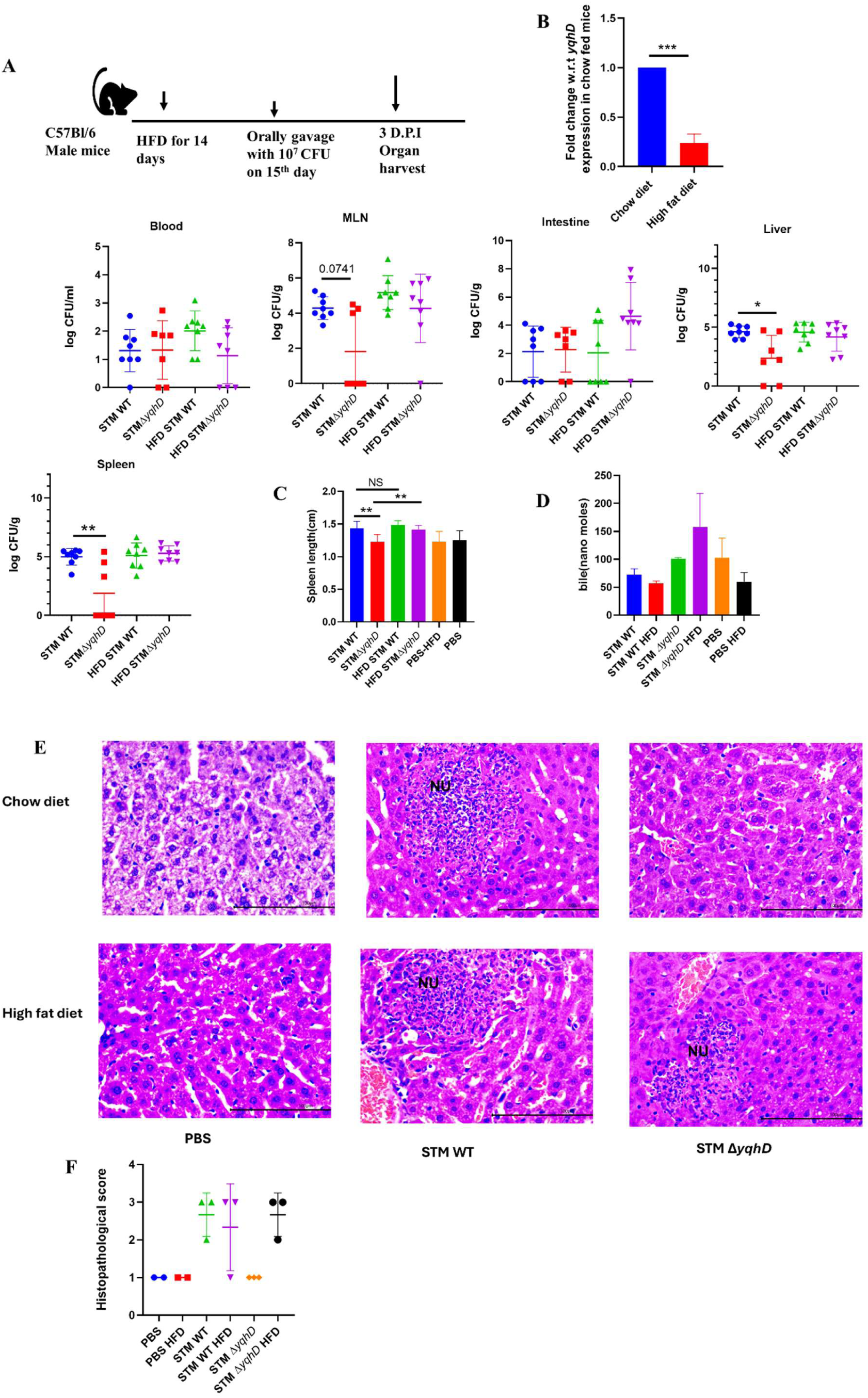
High-fat diet increases organ burden of STM Δ*yqhD* in secondary sites of infection in C57BL/6 mice. **A**. Schematic of infection and bacterial organ burden of STM WT and STM Δ *yqhD* in C57BL/6 male mice 3 days post-infection (8 animals per group). **B**. mRNA expression of *yqhD* in STM WT upon infection in C57BL/6 mice upon exposure to a high-fat diet compared to chow-fed mice. **C**. Measurement of spleen length from mice infected with STM WT and STM Δ *yqhD* chow diet and high-fat diet. Analysis performed by Mann-Whitney test.’ p values****<0.0001, ***<0.001, **<0.01, *<0.05. **D**. Measurement of bile in the serum samples of mice. **E.** Representative image of hematoxylin-eosin-stained liver sections upon *Salmonella* infection at 3^rd^ days post-infection at 40X. Scale bar=100 µM. Here, NU is neutrophil aggregation. **F**. Quantification of the data in E. Pathology score was assigned according to histological activity index as-Severe −4; Moderate −3; mild −2; Minor/minimum−1.

Similarly, five days post-infection, there was a reduced organ burden of STM *ΔyqhD* when mice were on a chow diet compared to STM WT. It was significantly increased when mice were fed HFD (Supplementary Fig.4A). There was increased weight reduction in STM *ΔyqhD* infected mice when they were kept on HFD compared to STM WT (Supplementary Fig.4B). Similar observations were validated by measuring the spleen length (Fig 2C).

Furthermore, we have also measured the bile concentration in the mice serum samples. In accordance with previous literature, we observed decreased serum bile in HFD mice infected with STM WT or PBS compared to chow diet control (*19*). Interestingly, we observed an increased bile concentration in STM *ΔyqhD* HFD mice compared to the chow diet control (Fig 2D). This increased circulatory bile might be responsible for the increased organ burden of STM *ΔyqhD* in the liver and spleen on HFD. Additionally, upon hematoxylin-eosin staining, we observed that liver sections had minor pathology in the mice infected with STM *ΔyqhD* on the chow diet compared to the STM WT, which has several areas with neutrophil aggregation. Similar damage scores for STM WT and STM *ΔyqhD* were observed in mice on HFD with various regions of neutrophil aggregation (Fig 2E, 2F, Supplementary Fig.5). Thus, treating mice with HFD leads to an increased organ burden of STM *ΔyqhD* compared to the chow diet. Recent reports have shown different immune responses in male and female rats (*20*). We reproduced our experiment in female mice and observed a similar trend. STM Δ*yqhD* has a lesser organ burden than STM WT when kept on a chow diet, and it became similar to STM WT when mice were maintained on HFD (Supplementary Fig.6A and B). Suggesting that increased colonisation of STM *ΔyqhD* in mice is independent of the sex of mice. In the *in vivo* pathogenesis model, the treatment of oleic acid, a primary digestive product of lard used in the HFD to mice, has been reported to induce bile synthesis(*18*). We observed no significant difference in the colonisation of STM Δ*yqhD* compared to STM WT in the cecum, ileum, and faeces on the treatment in C57Bl/6 mice (Supplementary Fig 7).

### 2. Oxidative stress is essential for the enhanced survival of STM *ΔyqhD*

In line with the antioxidant function of *yqhD*, we observed attenuated survival of the STM Δ*yqhD* in macrophages, as seen in our *in vitro* study in RAW 264.7 macrophages and peritoneal macrophages from C57BL/6 mice (Fig 3A). This attenuated proliferation in macrophages might account for the reduced organ burden of STM *ΔyqhD* in C57BL/6 mice. Macrophages are essential for the lifecycle of *Salmonella* in the host. During the infection cycle of *Salmonella*, the bacteria reside in the macrophages and utilise the immune cells to disseminate to secondary sites of infection (*21*). Macrophages present a hostile environment to the pathogen by generating ROS, RNS (Reactive nitrogen species), antimicrobial peptides and nutrient deprivation. ROS is generated through the NADPH-phagocytic oxidase as an immediate measure to combat the invading pathogen (*22*). STM *ΔyqhD* proliferation was higher in peritoneal macrophages from mice deficient in the NADPH phagocytic oxidase(*gp91^phox−/−^)* (Fig 3A). We also determined the intracellular ROS levels upon bile salt exposure using Dichlorofluorescein Diacetate (DCFDA). In the presence of 7% bile, STM *ΔyqhD* showed a higher intracellular ROS (Fig. 3C). Interestingly, the survival of STM *ΔyqhD* in the presence of bile salts is increased, even though it has higher ROS. In the literature, oxidative stress genes (catalase and superoxide dismutases) are downregulated on bile salt exposure (*23*).

**Fig. 3.**
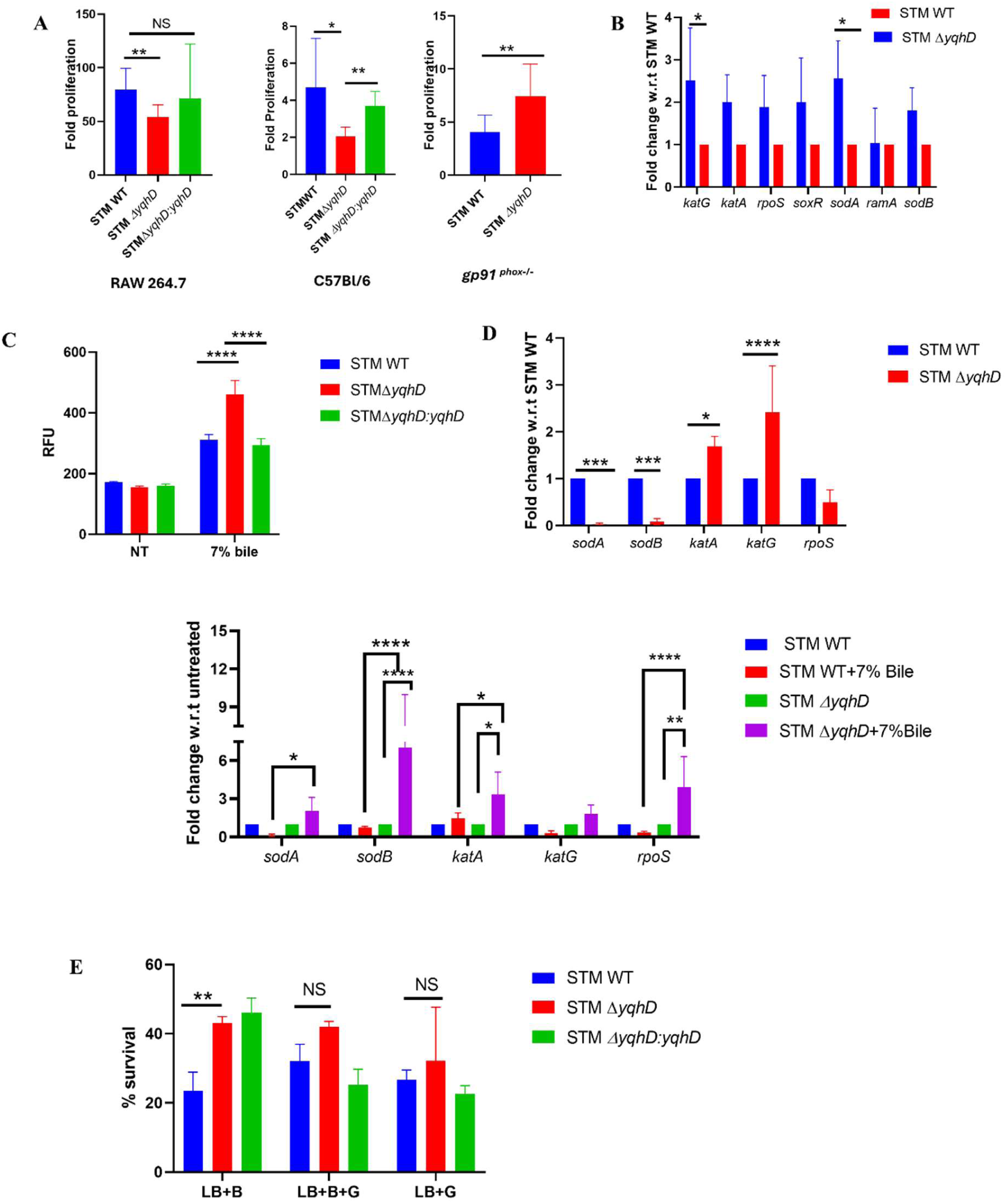
Antioxidative response of *yqhD* is essential for survival on bile salt exposure. **A.** Fold proliferation in RAW 264.7 macrophages and peritoneal macrophages from C57BL/6 and *gp91 ^phox^*^−/−^ mice of STM WT, STM Δ *yqhD* and STMΔ *yqhD:yqhD*. Analysis was performed using unpaired student’s t-test; p values****<0.0001, ***<0.001, **<0.01, *<0.05. **B**. The mRNA expression of antioxidant genes in STM Δ *yqhD* in RAW 264.7 macrophages at 16 hours post-infection. **C**. DCFDA assay to measure ROS levels in STM WT, STM Δ *yqhD* and STMΔ *yqhD:yqhD* on treatment with 7% bile. **D**. The mRNA expression of antioxidant genes in STM Δ *yqhD* as compared to STM WT and in the presence of 7% bile in the STM *ΔyqhD* and STM WT. **E**. The *in vitro* sensitivity assay on the treatment of STM *ΔyqhD* on exposure to bile salts in the presence of 10mM Glutathione by CFU/ml. Analysis was performed by using two-way ANOVA; p values****<0.0001, ***<0.001, **<0.01, *<0.05. All data are represented as mean±SD. Data is from one experiment representative of N ≥3 independent experiments.

**Fig. 4.**
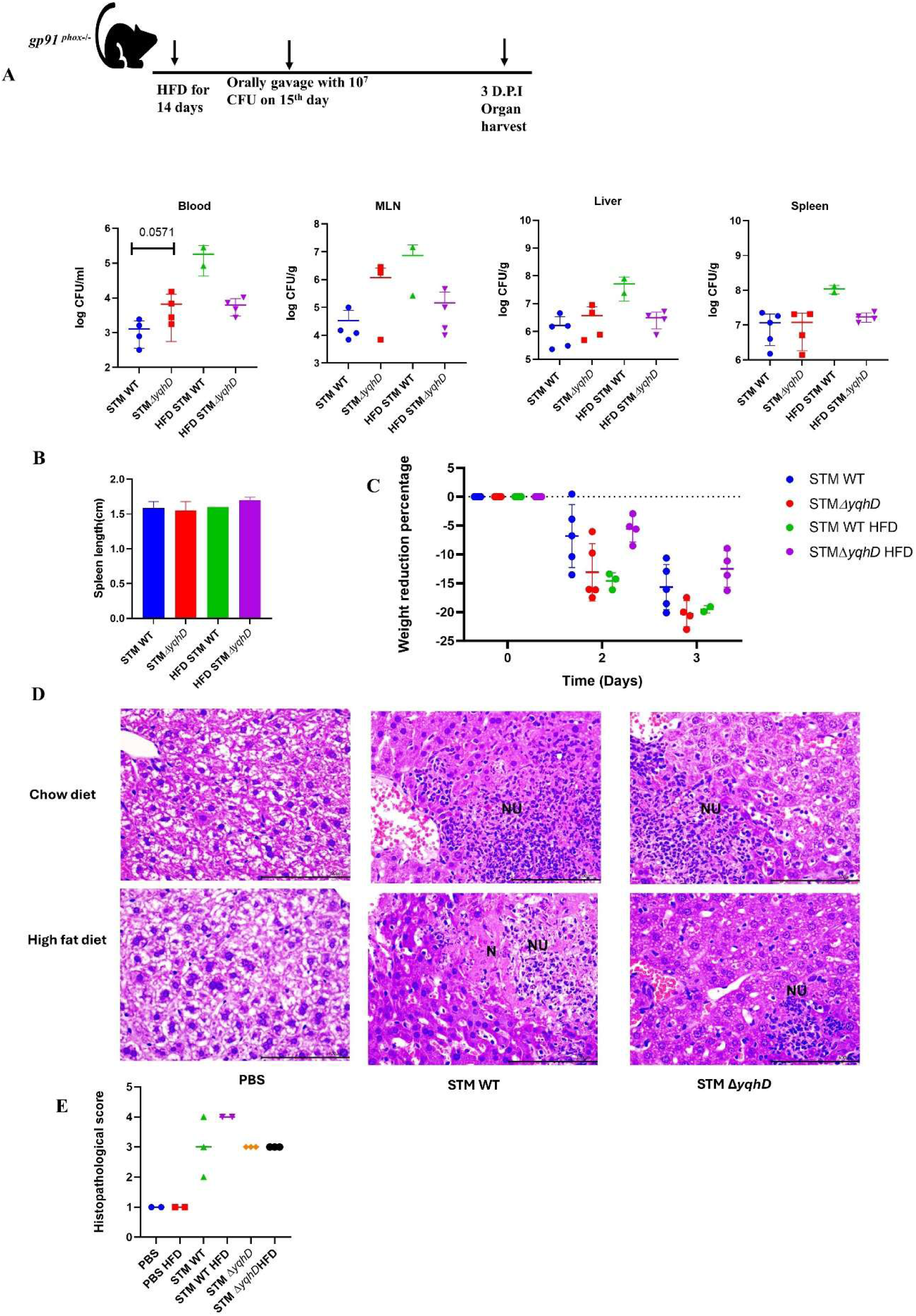
Oxidative stress is essential for increased organ burden of STM Δ*yqhD* in *gp91 ^phox−/−^*mice on treatment with a High-fat diet. **A.** Schematic of infection and organ burden of STM WT and STM Δ *yqhD* in *gp91 ^phox−/−^* - male mice 3 days post-infection (5 animals per group), Analysis performed by Mann-Whitney test. **B.** Average weight reduction on infection of STM WT and STM Δ *yqhD* in *gp91 ^phox−/−post−^*infection. **C**. Measurement of spleen length from mice infected with STM WT and STM Δ *yqhD* chow diet and high-fat diet. Analysis performed by Mann-Whitney test. **D**. Representative image of hematoxylin-eosin-stained liver sections upon *Salmonella* infection at 3^rd^ days post-infection at 40X. Scale bar=100 µM. Here, NU is neutrophil aggregation, and N is necrosis. **E**. Quantification of the data in D. Pathology score was assigned according to histological activity index as-Severe −4; Moderate −3; mild −2; Minor/minimum −1.

Similarly, in our results in STM WT, major oxidative stress genes are downregulated and upregulated in STM *ΔyqhD.* On qRT PCR analysis of various oxidative stress genes, we observed that STM *ΔyqhD* has upregulation of *katA, katG* and downregulation of *sodA, sodB.* On bile salt exposure, *sodA, sodB*, and stress response molecule *rpoS* were significantly upregulated. Thus, *sodA* and *sodB* are induced in the STM *ΔyqhD* in response to bile (Fig. 3D). In the HepG2 cells, we observed that all the antioxidant genes are downregulated in STM *ΔyqhD* except *soxR* (Supplementary Fig.8). To elucidate whether the increased survival of STM *ΔyqhD* was due to oxidative stress. Along with the primary bile salt treatment to STM *ΔyqhD,* antioxidant glutathione was added to the LB media. We observed that there was similar survival in STM WT and STM *ΔyqhD* (Fig. 3E). Taken together, these observations suggest that oxidative stress is essential for the increased survival of the STM *ΔyqhD* as compared to STM WT in bile salt exposure.

To determine the effect of oxidative stress and a high-fat diet, we used *gp91^phox−/−^* male mice and kept them on HFD for 14 days, and infection was given on the 15^th^ day. STM WT and STM *ΔyqhD* had a similar organ burden on the chow diet. At the same time, there was reduced organ burden in STM *ΔyqhD* as compared to STM WT on HFD (Fig. 3A). However, there was no difference in the spleen length (Fig. 3B). On the analysis of the weight post-infection, there was increased weight reduction in the mice infected with STM WT on HFD as compared to STM *ΔyqhD* (Fig.3C). Upon Haematoxylin and eosin staining of the liver samples from these mice we observed there was similar pathology in both STM WT and STM *ΔyqhD* on chow diet with neutrophil aggregation. While on HFD, there was severe pathology with STM WT infection having necrosis and neutrophil aggregation as compared to STM *ΔyqhD* (Fig. 3D, 3E and Supplementary Fig. 9). In conclusion, STM *ΔyqhD* requires oxidative stress for the increased organ burden on HFD, as seen in the C57BL/6 mice. In *gp91phox−/−* mice deficient in mounting an oxidative response, there is a comparatively reduced organ burden of STM *ΔyqhD* as compared to STM WT on HFD.

### 3. *yqhD* reduced the survival of *Salmonella* to bile by regulating the efflux pump activity

Upon exposure to bile, *Salmonella* modulates envelope structures like lipopolysaccharides, peptidoglycan (*ybiS*), and the enterobacterial common antigen (*wecD*) to provide a barrier against the bile salt uptake (*4, 24, 25*). Bile also downregulates porins (*ompF* and *ompC*) to reduce the uptake of bile salts (*6*). Furthermore, the efflux systems(*acrAB)* are induced, which decreases the intracellular concentration of bile salts (*26*). Exposure to bile also upregulates the DNA damage genes(*recA* and *dinP)* to overcome the damage caused by bile(*8*).

To elucidate the mechanism behind the increased survival of the *yqhD* knockout, we performed qRT PCR analysis for various bile stress genes reported in the literature for enteric bacteria. We observed a significant decrease in mRNA expression of the efflux pump repressor (*acrR*) and porins (*ompF* and o*mpC*) in the *yqhD* mutant upon treatment with bile salts (Fig. 5A). Similarly, upon infection into the HepG2 cells, the *yqhD* mutant executed a time-dependent decrease in the mRNA levels of efflux pump repressor and porins (Fig. 5B). To validate the regulation of efflux pump and porin activity by *yqhD* mutant further, we determined the efflux pump and porin activity using the Bis-benzimide assay. Bis-benzimide is a nuclear-binding dye; its fluorescence depends on porins and efflux pump activity. We found a decreased fluorescence for the *yqhD* mutant compared to STM WT. The decrease in fluorescence could be attributed to increased efflux pump activity or the low expression of the porins. So, if the efflux pump were mediating the reduction in fluorescence on treatment with a known efflux pump inhibitor CCCP(Carbonyl cyanide m-chlorophenyl hydrazone), it would lead to similar fluorescence levels in the wild type and the mutant, as observed (Fig. 5C). To further strengthen the observation that the efflux pump activity mediates the enhanced survival of the *yqhD* mutant, we generated an *acrB* mutant in the background of *yqhD* deletion. The STM*ΔacrBΔyqhD* showed no growth upon bile salt exposure compared to STM*ΔacrB,* which showed highly attenuated growth (Fig. 5D). Thus, the growth advantage of STM*ΔyqhD* under bile stress is lost upon deletion of both *acrB* and *yqhD*. Similarly, we observed a similar reversal in fold proliferation of STM*ΔacrBΔyqhD* upon infection into the HepG2 cells (Fig. 5E). Our results suggest that *yqhD* mediates sensitivity to bile by regulating the AcrAB efflux pump activity.

**FIG. 5.**
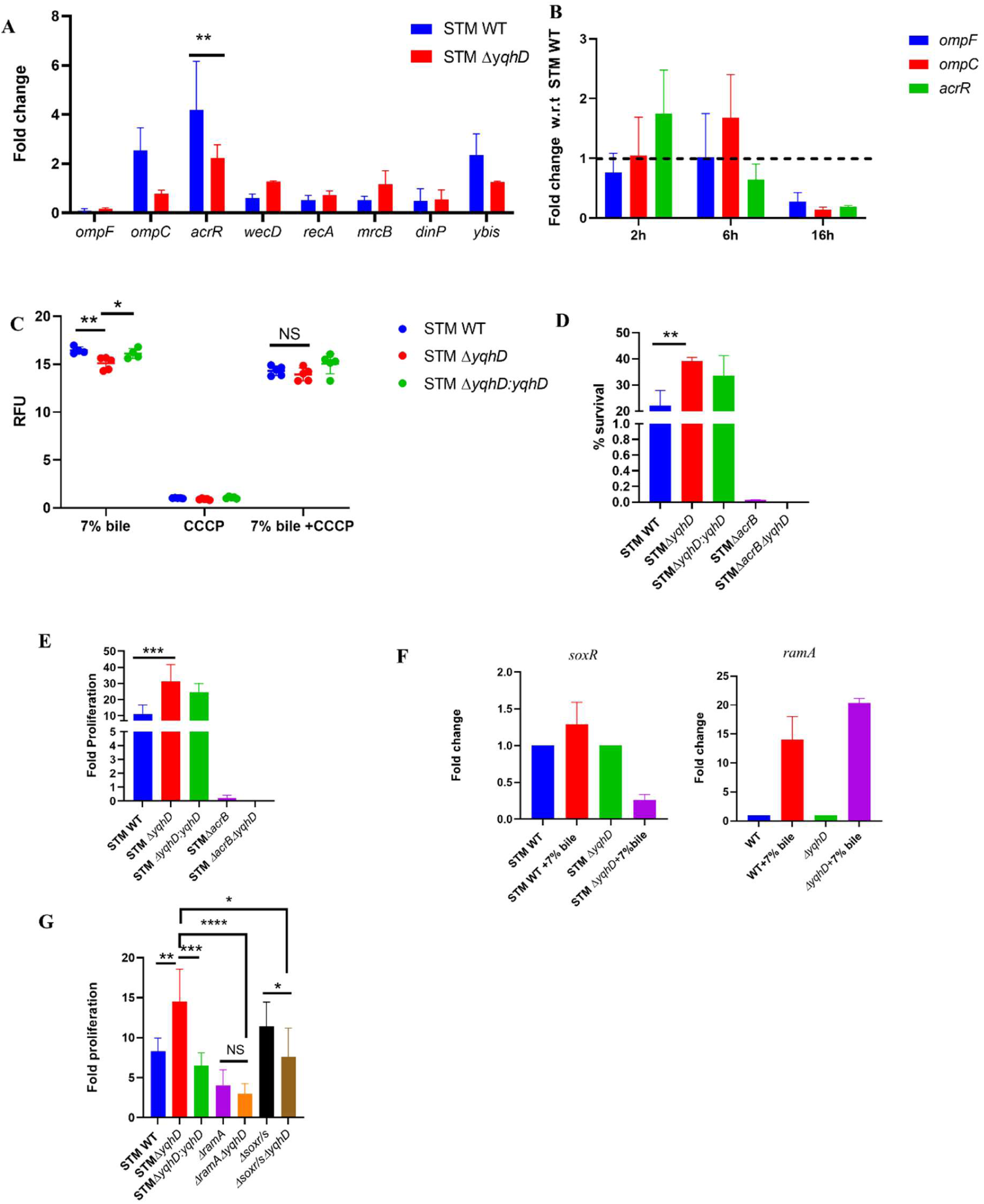
Deletion of *yqhD* downregulates AcrAB efflux pump repressor using RamA/R regulon. **A**. The mRNA expression of various bile-responsive genes in STM WT, STMΔ*yqhD* on exposure to bile salts in LB Media. **B**. The mRNA expression of porins and efflux pump repressor in HepG2 cells using qRT-PCR (N=3,n=3). **C**. Bis-benzimide assay to assess the efflux pump and porins activity in the presence of bile salt with and without efflux pump inhibitor CCCP (N=3,n=5), Analysis performed using two-way ANOVA, p values****<0.0001, ***<0.001, **<0.01, *<0.05. **D**. The *in vitro* sensitivity assay on treatment t of 7% bile salts to STM WT, STM *ΔyqhD,* STM *ΔyqhD:yqhD,* STM *ΔacrB,* STM *ΔacrB ΔyqhD* (N=3,n=2) in LB media. **E**. Fold proliferation of STM WT, STMΔ*yqhD* and, STMΔ*yqhD:yqhD,* STM *ΔacrB,* STM *ΔacrB ΔyqhD* in HepG2 cells (N=3,n=3). Analysis performed using unpaired Student’s t-Test.; p values****<0.0001, ***<0.001, **<0.01, *<0.05. **F**. The mRNA expression of *soxR and ramA* in STM *ΔyqhD* and STM WT on treatment with and without bile salts in LB media using qRT PCR(n=3, N=3) **G**. Fold proliferation of STM WT, STMΔ*yqhD* and, STMΔ*yqhD:yqhD,* STM *ΔacrB,* STM *ΔacrB ΔyqhD,* STM *ΔramA,* STM *ΔramAΔyqhD,* STM *ΔsoxR/S,* STM *ΔsoxR/SΔyqhD* in HepG2 cells (N=3,n=3). Analysis is performed using unpaired Student’s T test; p values****<0.0001, ***<0.001, **<0.01, *<0.05.

The fluoroquinolone susceptibility has also been reported to be highly dependent on the AcrAB efflux pump (*27*). We hypothesised that upon exposure to fluoroquinolones (ciprofloxacin), *yqhD* deletion would lead to antibiotic resistance. Moreover, pre-exposure to bile salts leads to adaptation and increases the MIC of antibiotics (*5*). To assess antibiotic susceptibility, we employed the micro-broth dilution method. We observed that the *yqhD* mutant was more susceptible than the wild type with and without pre-treatment with 7% bile (Supplementary Fig. 10A). Moreover, upon the treatment of bacteria with the MIC concentration (0.0078 µg/ml) of ciprofloxacin, no difference in the porosity and efflux pump activity was observed for the mutant compared to the wild type (supplementary Fig. 10B). Thus, our results conclude that *yqhD* plays a role in fluoroquinolone resistance. However, the mechanism for fluoroquinolone resistance differs from its action in the presence of bile.

### 4. *yqhD* regulates the efflux pump activity using the RamA/R two-component system

YqhD is a cytosolic protein, and the AcrAB efflux pump is located on the inner membrane of *Salmonella*. Hence, an interesting question arises: how does YqhD regulate the efflux pump activity? We hypothesised that elevated oxidative stress upon the deletion of *yqhD* leads to the activation of efflux pump activity.

It has been reported previously that RamA/R and SoxR/S regulons are activated under oxidative stress and lead to the induction of efflux pump. The AcrAB efflux pump is regulated by Mar operon in *E.coli*, while *ramA* is the major regulator of *Salmonella*, and *soxR/S* plays a small role in the induction (*28*). RamA acts on the *acrAB* efflux pump by binding to bile salts (cholic acid). RamA is transcriptionally repressed by RamR (*29–31*). RamA was also reported to bind directly to alcohol dehydrogenase P(*aldhP*) in *Klebsiella pneumoniae* (*32*). Another study showed that paraquat, a superoxide generator, could induce the *acrAB* efflux pump using SoxS, independent of RamA (*33*). Upon treatment with bile salts, we assessed the mRNA levels of both *ramA* and *soxR* and observed an increase in *ramA* and a decrease in *soxR* expression in the *yqhD* mutant compared to STM WT (Fig. 5F). We further generated deletion mutants for *ramA* and *soxR/S* regulon in the background of the *yqhD* mutant. We observed that STM *ΔramAΔyqhD* showed reduced survival upon exposure to 7% bile salts as compared to STM *ΔyqhD* and STM *ΔramA* measured by spotting on LB-agar plate supplemented with 7% bile salt. While *ΔsoxR/SΔyqhD* showed enhanced survival than STM *ΔsoxR/S* (Supplementary Fig. 11). Similarly, in HepG2 cells, the proliferation of the *yqhD* mutant was significantly reduced when *ramA* was mutated. The single deletion of *ramA* has also attenuated the proliferation in HepG2 cells (Fig. 5G). These observations lead us to conclude that RamA/R regulon mediates the increased survival of STM *ΔyqhD* upon bile salts exposure and is essential for survival in HepG2 cells.

### 5. Bile stress doesn’t reduce invasion in STM Δ*yqhD*

Our study suggests that the absence of YqhD in *Salmonella* is advantageous for its survival in the presence of bile. Thus, we wanted to question how *Salmonella* utilises YqhD to its advantage. Interestingly, we observed that the invasion in hepatocytes (HepG2) was significantly reduced in STM Δ*yqhD* as compared to STM WT (Supplementary Fig. 12B). While the invasion into the colon carcinoma cells (Caco−2) was similar in STM WT and STM Δ*yqhD* (Supplementary Fig. 12A). These results indicate that the invasion of the mutant is altered in the in HepG2, unlike Caco−2 cells. The reduction of invasion in HepG2 cells could be due to the downregulation of SPI−1 machinery by bile salt produced by Hepatocytes.

*Salmonella* invasion in eukaryotic cells is mediated by Type three secretion system 1 encoded by *Salmonella* pathogenicity island 1(SPI−1). Bile is an environmental cue to repress SPI−1 genes by destabilising the HilD (*34–36*). SPI−1 genes are also repressed by *ramA*, which is expressed by exposure to bile (*37*). In our study, we also observed that *yqhD* is regulated by *ramA.* To understand whether the reduced invasion of STM Δ*yqhD* is due to the modulatory function of bile in the expression of SPI−1 genes, we treated STM WT and STM Δ*yqhD* with 7% bile salt in LB media. Compared to the untreated samples, we observed downregulation of the SPI−1 encoded genes (*invF, hilA, sipA*) in STM *ΔyqhD*. Meanwhile, in STM WT, only *invF* was downregulated, and *sipA* and *hilA* were slightly upregulated (Supplementary Fig. 12 C). Thus, to further confirm our results, we observed the invasion with the knockdown of the CYP7A1 in HepG2 cells. Still, it could not rescue the invasion of STM *ΔyqhD* (Supplementary Fig. 12D). Thus, our results suggest that only bile secretion is not responsible for the reduced invasion of STM *ΔyqhD* in the HepG2 cells. There might be other factors responsible.

## Discussion

In modern times, the development of the fast-food industry has changed human lifestyle, and the high-fat diet (HFD)/Western diet has become popular. The Western diet contains 36 to 40 per cent fat(39). An HFD induces inflammation and dysbiosis of the gut health by increasing the ratio of *Firmicutes* to *Bacteroidetes.* It also causes the growth of *Enterobacteriaceae* (*38, 39*). HFD induces bile production, aiding the survival of bile-resistant bacteria like *Bilophila wadsworthia* (*40*) and killing bile-sensitive bacteria. *Salmonella* is also a bile-resistant pathogen, and bile salt production by HFD provides an ambient condition for its growth(*18*). This study describes the growth of *Salmonella* Typhimurium 14028S and Typhi CT18 on exposure to a bile salts mixture in vitro and liver carcinoma (HepG2 cells), as the liver is responsible for bile production. Bile salts are antimicrobial and dissolve membrane lipids, causing DNA damage and protein misfolding. Enteric bacteria have strategies to cope with bile-induced damage. Enteric bacteria modify their LPS structure, and long O Antigen increases the bile resistance by modulating the envelope (*41*). Bile salts that enter are also effluxed out using an AcrAB efflux pump (*7*). Bile salts-induced DNA damage is also repaired by homologous recombination and SOS-associated DNA repair (*8*). All the bile resistance in laboratory conditions has been studied by adding ox bile or individual bile salt like sodium deoxycholate to the bacterial media. Despite the significant number of studies, the genes involved in bile resistance keep increasing.

Our study addresses the novel interaction between the oxidative stress gene *yqhD* and bile salts. YqhD is studied widely in *E.coli* for its antioxidant function and importance in biofuel production. In addition to its role as an antioxidant, we found that YqhD increased susceptibility to Bile. Treatment of mice with sodium cholate and HFD led to increased colonisation by STM Δ*yqhD.* In the earlier studies, the role of major antioxidant genes on bile salt treatment is reported to be negligible except for disulfide stress, and their deletion does not confer any disadvantage to the bacteria (*42*). The increased survival of the *ΔyqhD* was due to increased oxidative stress on the exposure to bile salts, and treatment with Glutathione *in vitro* and deletion of *gp91^phox−/−^* in mice reversed the advantage.

AcrAB efflux pump is widely studied; it provides resistance against decanoate, bile salts, and ethanol (*43*). AcrAB efflux pump has also been reported for its role in colonisation, persistence, and invasion (*44*). In our study, *yqhD* induced the activity of the AcrAB efflux pump and deleting the *acrB* gene of the efflux pump led to reduced survival of STMΔ*yqhD* on bile salt exposure. RamA has been reported in the literature to respond to bile stress and induce AcrAB efflux pump (*29, 30*). AcrAB efflux pump was modulated by YqhD using RamA/R regulon. The current study has limitations regarding how exactly *yqhD* modulates the RamA/R regulon. Despite the improved survival on deletion of *yqhD* in the presence of bile salt, there is a significant reduction in invasion due to the downregulation of SPI−1. STM Δ*yqhD* survival on bile salt exposure was found to be mediated by RamA/R regulon, which also can mediate the invasion of *Salmonella* (*37*). However, we observed that knockdown of the bile salt synthesis could not rescue the invasion, thus suggesting the invasion of STM Δ*yqhD* is independent of Bile. Therefore, from our study, we conclude that YqhD is both beneficial and detrimental to S*almonella.* The mechanism of reduced invasion of STM Δ*yqhD* in hepatocytes remains yet to be answered.

This study aims to shed light on the survival of *Salmonella* when exposed to bile salts. Knowledge of the mechanism of bile resistance by *Salmonella* and other food-borne pathogens may stimulate novel tools to combat diseases. If the *Salmonella* burden in the gall bladder of chronic carriers could be reduced, it would reduce transmission of the disease and the burden on public healthcare.

## Acknowledgements

Dr Abhilash Vijay Nair and Debapriya Mukherjee have helped in the oral gavage of the animals. We sincerely thank them. We acknowledge the Departmental Real-Time Facility and Central Animal facility at IISc for supporting the experiments.

## Author contributions

Conceptualisation-KP and DC

Methodology-KP and DC

Investigation-KP and YK

Analysis-KP

Writing and Editing -KP and DC

## Data and material availability

All data are available in the main text or supplementary materials.

## Material and methods

1. Bacteria culture *-Salmonella enterica* serovar Typhimurium 14028S strain used in the study was a kind gift from Prof. Michael Hensel. The bacterial strains were revived from glycerol stocks at −80°C and streaked on the LB agar plates with the appropriate antibiotics. For the majority of bacterial cultures, a single colony was taken and inoculated in an LB tube and grown at 37°C, 170rpm, in a shaking incubator. STM pkD46 was grown at 30°C, 170rpm in the shaking incubator.

The growth curve was performed by subculturing(1:100) the overnight culture in Fresh Lb media, and growth was monitored for 24 hours using Bioscreen.

| S.NO | Bacterial strain | Antibiotic |
| --- | --- | --- |
| 1 | *Salmonella* Typhimurium 14028S (STM WT) | - |
| 2 | STM $\Delta yqhD$ | kanamycin |
| 3 | STM $\Delta yqhD:yqhD$ | Kanamycin, ampicillin |
| 4 | STM $\Delta acrB$ | Chloramphenicol |
| 5 | STM $\Delta acrB \Delta yqhD$ | Kanamycin, Chloramphenicol |
| 6 | STM $\Delta ramA$ | Chloramphenicol |
| 7 | STM $\Delta ramA \Delta yqhD$ | Kanamycin, Chloramphenicol |
| 8 | STM $\Delta soxR/S$ | Chloramphenicol |
| 9 | STM ΔsoxR/S ΔyqhD | Kanamycin, Chloramphenicol |
| 10 | *Salmonella* Typhi CT 18(STY) | - |
| 11 | STY ΔyqhD | Kanamycin |

2. Knockout and complement generation-Knockout strains were made using the one-step gene inactivation strategy used by Datsenko and Wanner (*16*). STM WT with pKD46 plasmid, having lambda red recombinase system under arabinose inducible promoter, was grown in LB media supplemented with 50µg/ml ampicillin and 50mM arabinose at 30°C. Bacterial culture was sub-cultured (1:33) and incubated for 3 hours to achieve log phase culture. Electrocompetent STM pKD46 cells were prepared by washing the bacterial cells with Milli Q water and 10% glycerol. Kanamycin and chloramphenicol resistance gene cassettes amplified from pkD4 and pkD3 using gene-specific knockout primers were used for the electroporation. The transformed bacteria were selected on LB agar with appropriate antibiotics. Transformed bacterial colonies were patched on the fresh antibiotic plate and confirmed with gene-specific confirmatory primers.

Construction of the complement strain - *yqhD* gene was amplified by cloning primers using colony PCR. The PCR product and pQE60 plasmid were digested with restriction enzymes – EcoRI(NEB) and HindIII (NEB) in a cut smart buffer for 2 hours at 37°C. Digested insert (PCR product) and vector (pQE60) were gel purified and ligated by T4 DNA ligase(NEB) in 10X ligation buffer overnight at 16°C. The recombinant plasmid was transformed into the STM *ΔyqhD.* The recombinant plasmid was further confirmed by restriction digestion.

Construction of double knockout strains-single knockout strains of *ramA*, soxr/s and *acrB* were transformed with pKD46 plasmid. Transformed bacteria were grown with ampicillin(50µg/ml), chloramphenicol(25 µg/ml) and arabinose(50 mM) to log phase culture. The Kanamycin resistance gene cassette amplified from pKD4 plasmid was electroporated. The double knockout strains were selected on an LB-agar plate with kanamycin(50µg/ml) and chloramphenicol(25 µg/ml). Confirmatory primers further confirmed double knockouts.

3. Oxidative stress experiments-reactive oxygen species(ROS) in the bacteria were measured using redox-sensitive probe 2’,7’ dichlorodihydrofluorescein diacetate (H2DCFDA). H2DCFDA, once internalised, is oxidised by intracellular esterase and converted to fluorescent 2’, 7’-Dichlorofluorescein(H2DCF) with excitation at 495nm and emission at 529nm. Bacteria OD was adjusted to 0.3, subjected to bile stress overnight in LB, and incubated with 10µM of DCFDA for 30min. Fluorescence was measured.

4. Survival on bile exposure-log phase culture of the bacteria was adjusted to 0.3 OD in 1ml of phosphate buffer saline(PBS). Approximately 10^5^ bacteria were transferred to different concentrations of bile salt mixture (himedia) in Luria Broth. Treated bacteria were transferred to 96 well plates and incubated for 18 hours in the shaking incubator at 170rpm, 37°C. Post-incubation, absorbance was measured in a tecan plate reader at 595nm, and CFU analysis was performed by plating on LB-agar.

5. Bis benzamide assay-For measurement of membrane porosity and efflux pump activity. Bis benzamide dye(Sigma-Aldrich) was used with certain modifications to the previous protocol (*45*). Bacterial strains were grown overnight in LB broth, and absorbance was adjusted to 0.1 in 1X PBS. 7% bile alone or combined with cyanide 3-chlorophenylhydrazone (CCCP) was added to the bacterial cultures. 20µl of bisbenzimide was added to the 180 µl of bacterial culture in the 96-well plate and incubated for 10 minutes in the shaking incubator. Fluorescence was measured using a Tecan plate reader using a 346nm excitation and 460nm emission.

6. Q-RT PCR analysis-RNA-CDNA from bacteria and cell culture-the bacterial cells were pelleted down at 6000rpm for 10min. Pellets were lysed with TRIzol reagent (RNAiso plus) and stored at −80°C overnight. The lysed supernatant was subjected to chloroform extraction and precipitation with an equal volume of isopropanol. The precipitated RNA was air-dried and dissolved in DEPC-treated water. RNA concentration was checked using nanodrop, and the quality of the RNA was assessed through 1% agarose gel electrophoresis. cDNA was prepared post-DNase treatment (Thermo Fisher Scientific) using a PrimeScript RT reagent kit from Takara. Q-RT PCR was done using TB green in the bio-rad real-time detection system.

7. Cell culture – RAW 264.7 and HepG2 cells were maintained in DMEM(Lonza) supplemented with 10% FBS(Gibco) at 37°C in 5% CO2. Peritoneal macrophages were maintained in RPMI (Lonza) with 10 %FBS and 1% penicillin streptomycin(Sigma-Aldrich).

8. Isolation of peritoneal macrophages-Peritoneal macrophages were isolated from C57BL/6 mice and *gp91phox* knockout mice using a previously standardised protocol (*46, 47*). The mice were intraperitoneally injected with 8% Brewer’s Thioglycolate (from HiMedia). Five days post-injection, the primary macrophages were isolated from the peritoneal cavity. Any residual erythrocytes were lysed using an RBC lysis buffer (Sigma-R7757), and the isolated cells were maintained in an RPMI 1640 medium with 10% FBS and 0.1% penicillin-streptomycin for further experiments.

9. Virus-mediated knockdown of CYP7A1 in HepG2 cells-shRNA constructs with puromycin as a selection marker was a kind gift from Professor Subba Rao. Stable cells with CYP7A1 knockdown were prepared in HepG2 cells by lentivirus transduction, then gradually selecting the cells with puromycin. The cells were maintained in media containing 1 µg/ml puromycin. Lentivirus using HEK293T cells is prepared following a previously described protocol (*48*).

shRNA1- CCGGCCACAGTTAATGCACTTAGATCTCGAGATCTAAGTGCATTAACTGTGGTTTTTG

shRNA2- CCGGCCACCTCTAGAGAATGGATTACTCGAGTAATCCATTCTCTAGAGGTGGTTTTTG

10. Intracellular survival assay- 0.2 million RAW 264.7 were infected with stationary phase culture of the different bacterial strains. The multiplicity of infection(MOI) used for the RAW 264.7 was 10. HepG2 cells were infected with log phase culture and MOI of 10. For RNA isolation from the HepG2 cells, 1 million cells were infected with an MOI of 25 and lysed with Trizol at different time points.

Briefly after the infection with bacteria, the cells were centrifuged for 5 min at 800 rpm, followed by incubation at 37°C in 5% CO2 for 25 min. Cells were washed with 1X PBS and incubated with complete media containing 100µg/ml of gentamycin to kill the bacteria outside the eukaryotic cells for 1 hour. Subsequently, cells were washed with 1X PBS and complete media containing 25 µg/ml of gentamycin was added to prevent any secondary infection. Cells were lysed post 2 hours of 100µg/ml of gentamycin using 0.1 % triton X 100 to calculate per cent phagocytosis and per cent invasion. The bacterial number was determined by plating on LB agar. Similarly, the bacteria used in the infection was determined by plating the pre-inoculum. The ratio of bacteria at 2 hours in cell lysate to pre-inoculum multiplied by 100 gave us per cent phagocytosis for macrophages and per cent invasion for the epithelial cells. For calculations of fold proliferation, cells were lysed at 16 hours and 2 hours post-infection. The bacterial number was determined by plating on LB-agar. Fold proliferation is the ratio of the bacterial CFU at 16 hours to bacterial CFU at 2 hours.

11. Animal experiments- 5–6-week-old C57BL/6 or *gp91^phox−/−^* mice were used for the *in vivo* infection and peritoneal macrophage isolation. The institutional animal ethics committee approved all the experiments. The ethical clearance number for this study is CAF/Ethics/852/2021 and CAF/Ethics/021/2023.

10mg/ml oleic acid and 8% sodium cholate were orally gavage to 5–6-week-old C57BL/6 mice one hour before infection and four hours after infection. Oleic acid was dissolved in DMSO, and sodium cholate was dissolved in MQ. For both compounds, appropriate vehicle controls were used. Mice were infected with 10^7^ CFU/ml of STM WT, STM *ΔyqhD.* Mice were euthanised 24 hours post-infection, and ileum, cecum and faeces were collected.

Mice were fed a high-fat diet (60% from ICMR) for 14 days, and their weight was monitored and were infected with 10^7^ CFU/ml of STM WT, STM *ΔyqhD* in cohorts of five by oral gavage. On the third day, post-infection mice were sacrificed, and MLN, liver, and spleen were harvested for the bacterial burden, 3.5% Paraformaldehyde, before sample preparation for Haematoxylin and eosin staining. Isolated organs were homogenised and plated in *Salmonella Shigella* agar. All the CFU/ml obtained was normalised to the organ weight of the sample.

For histopathological scoring of hepatic necroinflammation/ pathology score: The comparison of pathological changes in the liver tissues was evaluated under light_microscopy by a veterinary pathologist and scored using a scientific method. The hepatic necroinflammation score is done according to the histological activity index (HAI) criteria with little modification(*49*). The scoring of necroinflammation is graded as 0−4 for each of the pathological lesions (Severe −4; Moderate −3; mild −2; Minor/minimum −1; No pathology −0) considering the following pathological lesions focal inflammation, focal necrosis, and portal inflammation/ Portal lymphohistiocytic.

12. Bile measurement-the experiment was performed per the kit protocol(ab239702). Briefly, serum samples harvested from the mice and stored at −80°C were thawed and added to the 96 well plate. The probe and enzyme mix were added, and absorbance was measured at 405 nm.

13. Determination of Minimum inhibitory concentration(MIC)−10^6^ bacteria of log phase culture of different bacterial strains treated with or without 7% bile were transferred to 96 well plates with varying concentrations of ciprofloxacin in Muller-Hinton broth (*50*). The bacteria were incubated for 18 hours in a shaking incubator at 37°C and 170rpm. After incubation, the MIC was determined by measuring absorbance at 595nm in a Tecan plate reader.

14. Spot assay-log phase culture bacteria adjusted to 0.3 OD was diluted and were used to spot 5µl on the LB agar plates with various concentrations of bile salt mixture or LB-agar alone. Plates were incubated overnight at 37°C. Images were acquired using Gel Doc.

15. Statistical analysis-GraphPad Prism 8.4.3 was used to perform all the statistical analysis. Analysis was done using unpaired two-tailed student t-tests and two-way ANOVA. In the animal experiments, the Mann-Whitney U test was performed.

---

**Supplementary Fig. 1.**
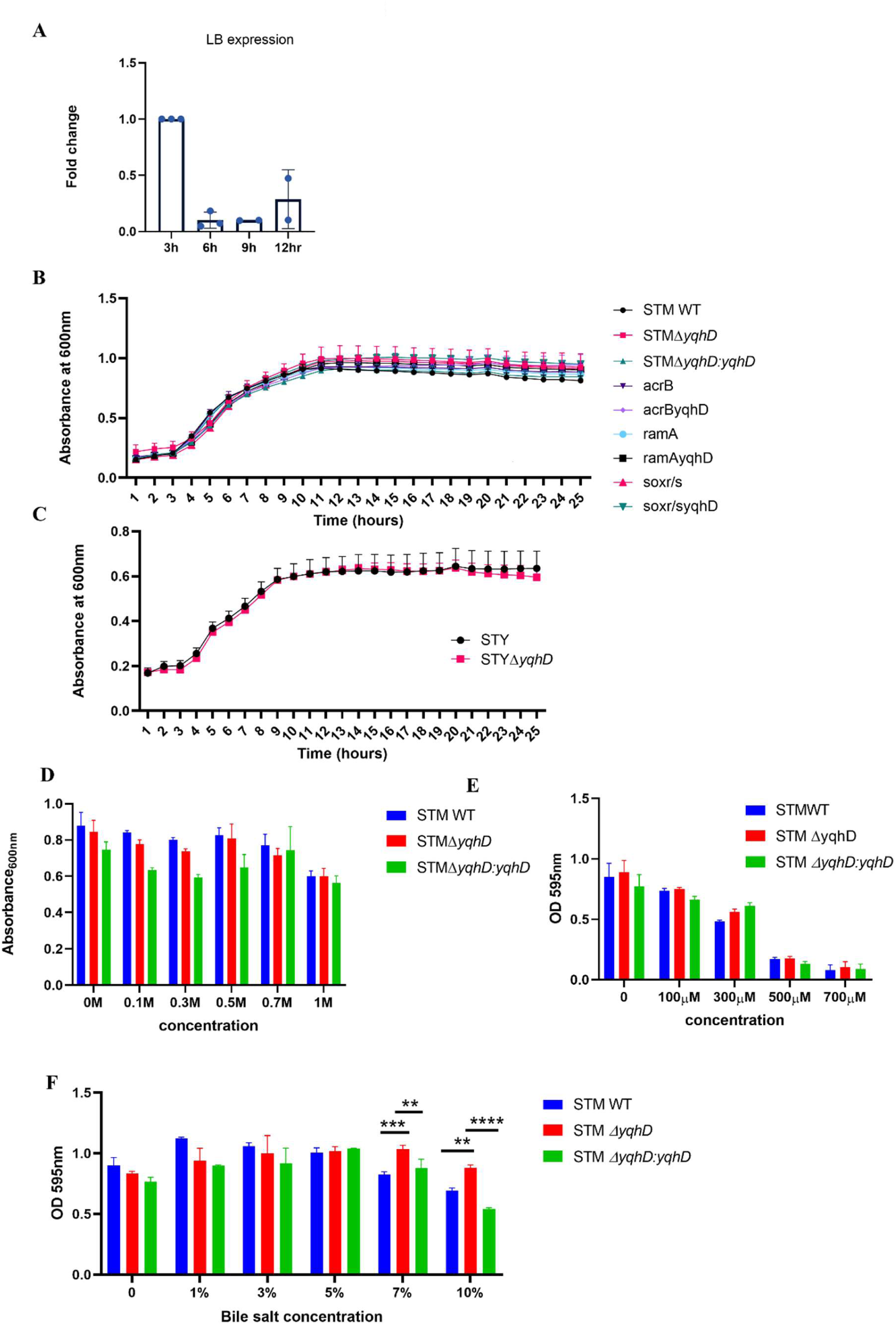
Growth kinetics of the various strains used in the study and their response to various stresses. **A**. The mRNA expression of *yqhD* at different time points in LB media in STM WT with respect to 3 hours. **B, C**. Growth kinetics of various strains used in the study. **D**. Absorbance at various concentrations of sodium chloride (N=3,n=3). **E**. Absorbance at various concentrations of iron chelator 2,2′-bipyridine.(N=3,n=3). **F**. Absorbance at various concentrations of bile salts (N=3,n=3). Analysis was performed using Two-way ANOVA; p values****<0.0001, ***<0.001, **<0.01, *<0.05..

**Supplementary Fig. 2.**
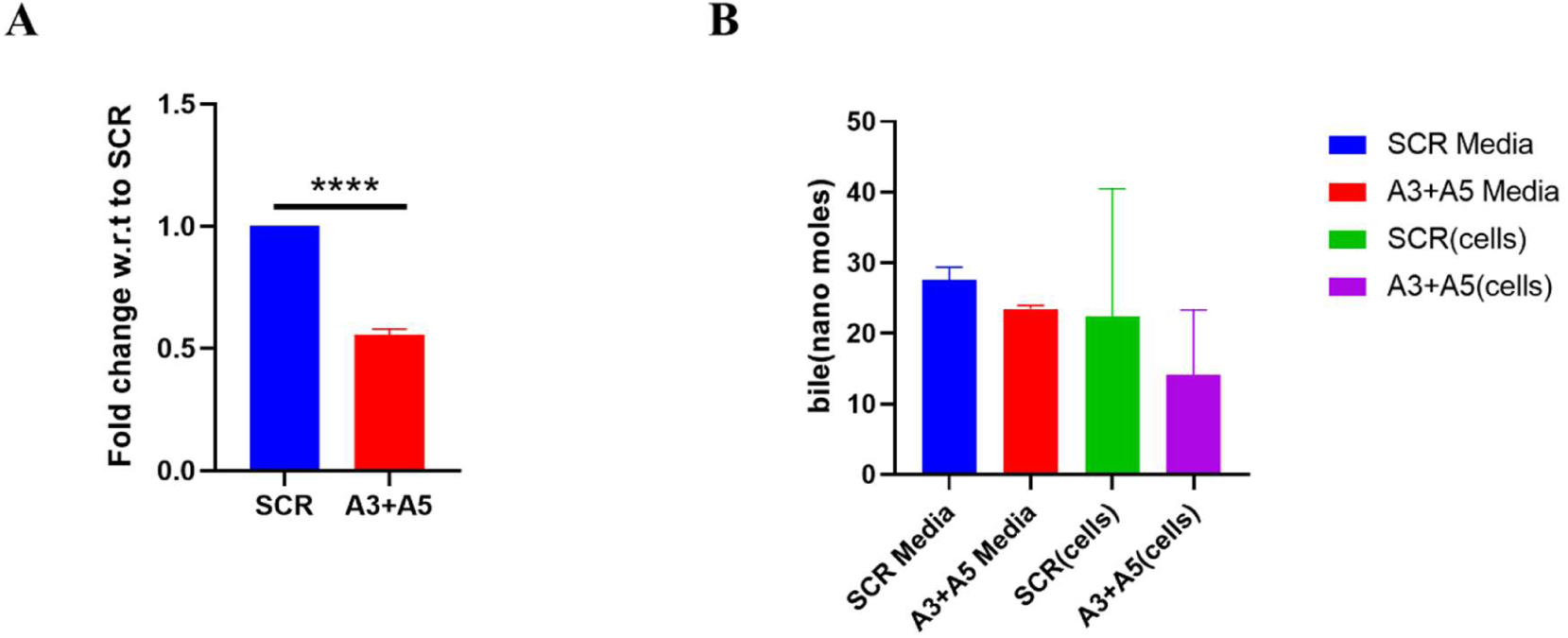
Knockdown of CYP7A1 in HepG2 cells. **A**. Validation of knockdown of CYP7A1 in HepG2 cells using qRT PCR. Analysis was performed using an unpaired two-tailed Student’s t-test. p values****<0.0001, ***<0.001, **<0.01, *<0.05. **B.** Bile estimation upon knockdown in HepG2 cells

**Supplementary Fig. 3.**
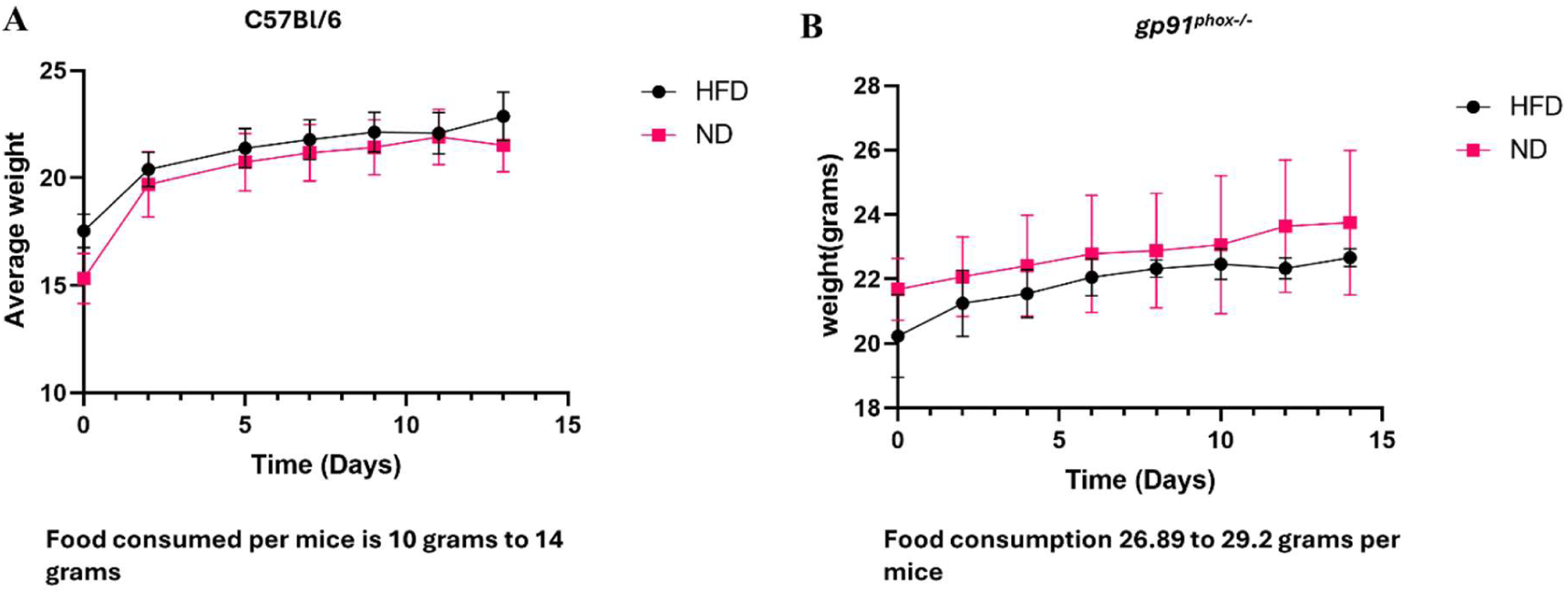
Average weight of mice fed on chow diet and High-fat diet. Average weight of mice on treatment with different diets for 15 days.

**Supplementary Fig. 4.**
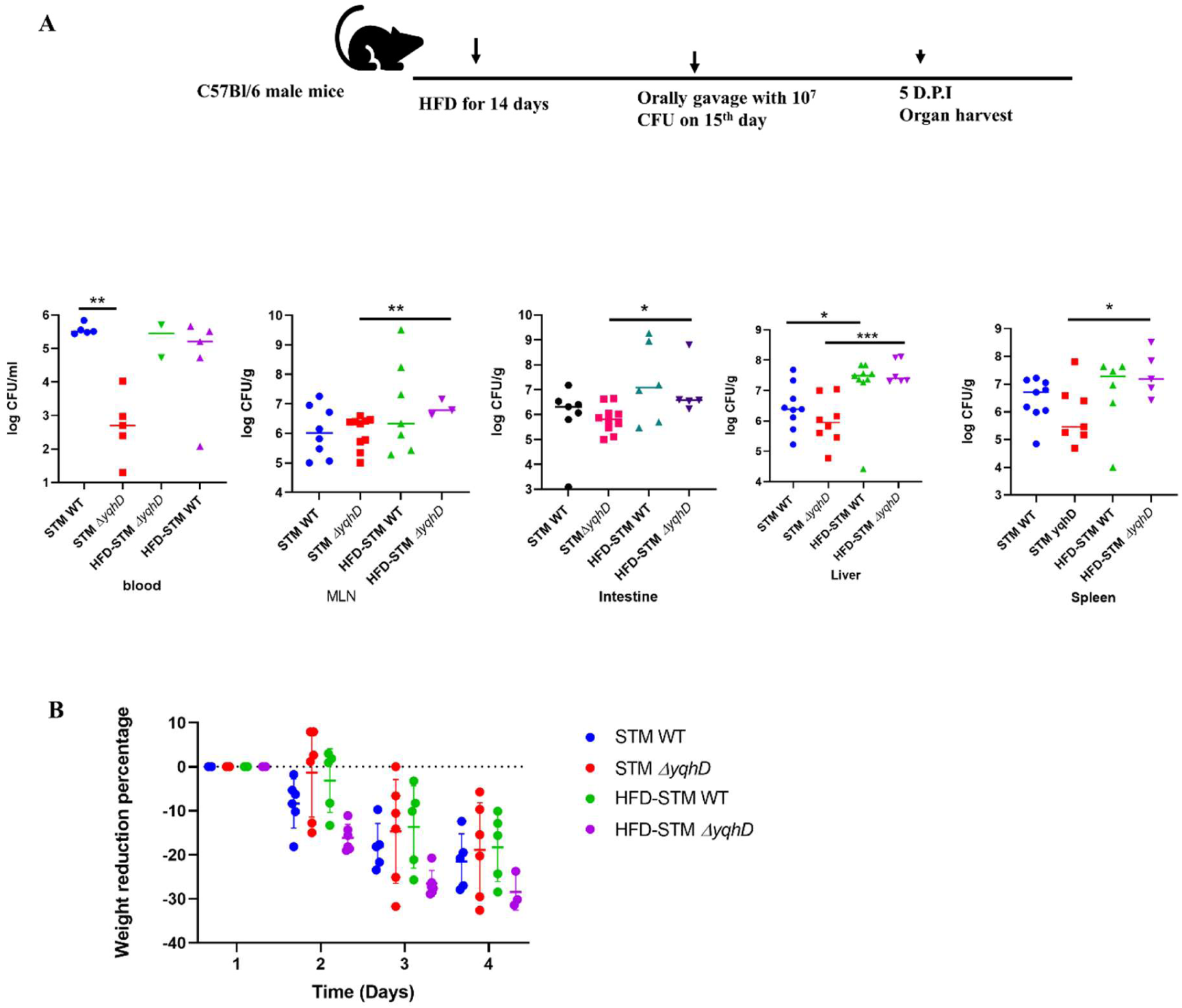
High-fat diet increases organ burden of STM Δ*yqhD* in C57BL/6 male mice at 5 days post-infection. **A.** Schematic of infection and organ burden of STM WT and STM Δ *yqhD* in C57BL/6 female mice 5 days post-infection, Analysis performed by Mann-Whitney test: p values****<0.0001, ***<0.001, **<0.01, *<0.05. Both dilutions have been used for plotting the data as there was mortality in STM *ΔyqhD* on the HFD treatment. **B.** Weight reduction after infection

**Supplementary Fig. 5.**
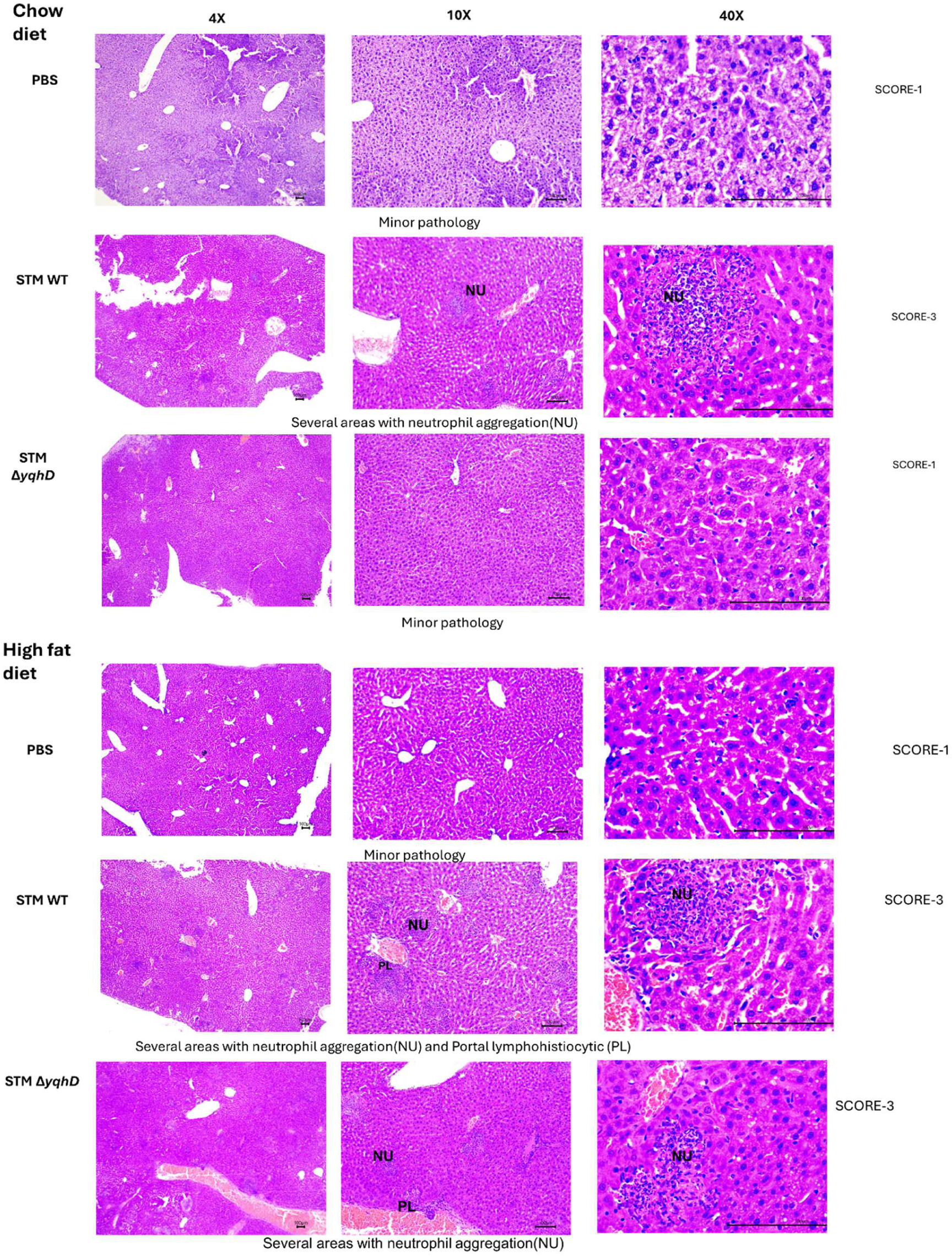
Haematoxylin and eosin staining of the liver sections from C57BL/6 mice at various magnifications with the pathology score. Histopathology images of liver sections used in the main figure 2 at different magnifications with their pathology scores.

**Supplementary Fig 6.**
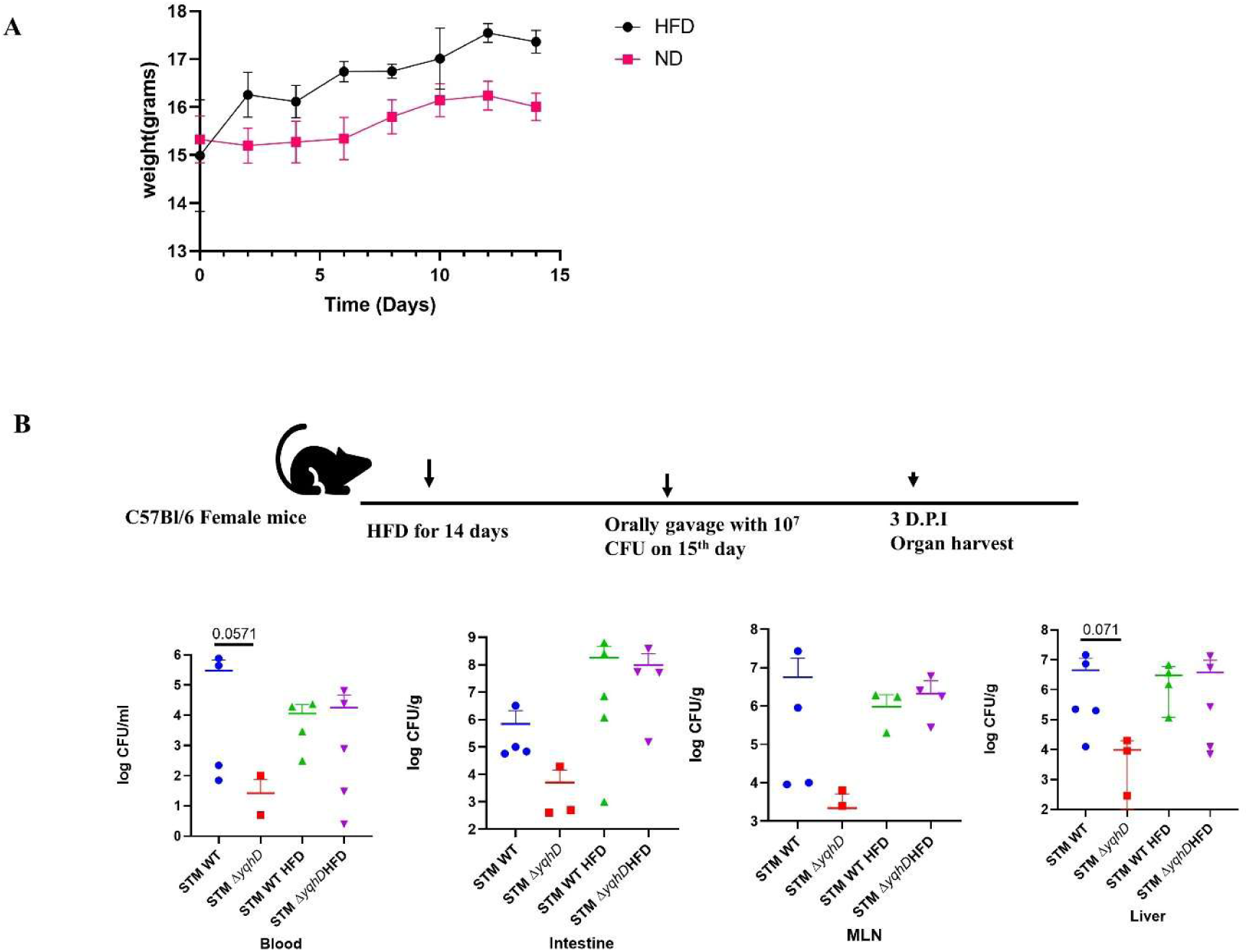
High-fat diet increases organ burden of STM Δ*yqhD* in C57BL/6 female mice. **A**. Average mice weight on treatment with HFD compared to chow diet (ND) **B**.Schematic of infection and organ burden of STM WT and STM Δ *yqhD* in C57BL/6 female mice 3 days post-infection, Analysis performed by Mann-Whitney test

**Supplementary Fig. 7.**
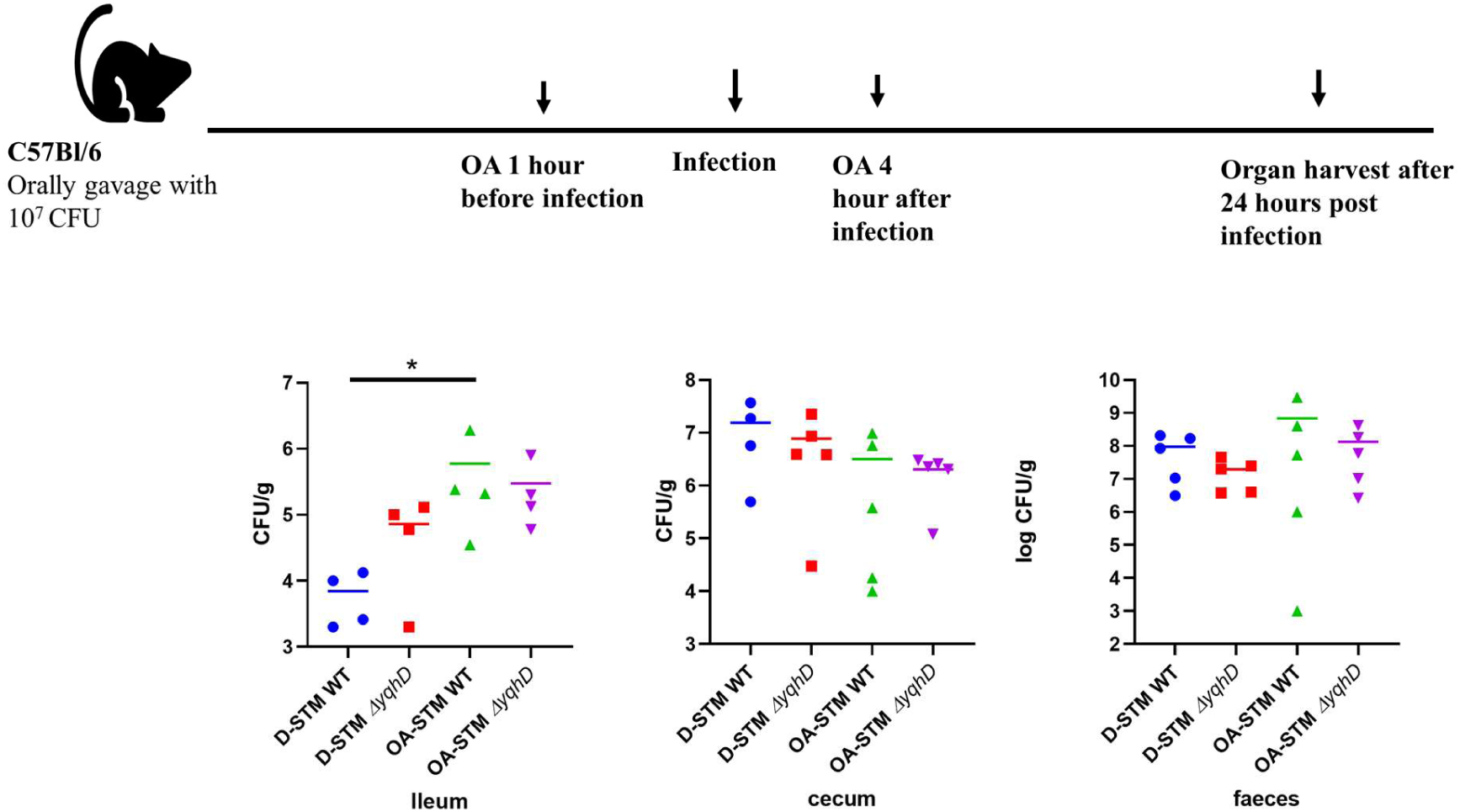
Oleic acid treatment and *Salmonella* colonisation in C57BL/6 mice. Schematic representation of animal infection and bacterial organ burden of STM WT, STMΔ*yqhD* in the ileum, Cecum and Faeces. Analysis by Mann-Whitney test. p values****<0.0001, ***<0.001, **<0.01, *<0.05.

**Supplementary Fig. 8.**
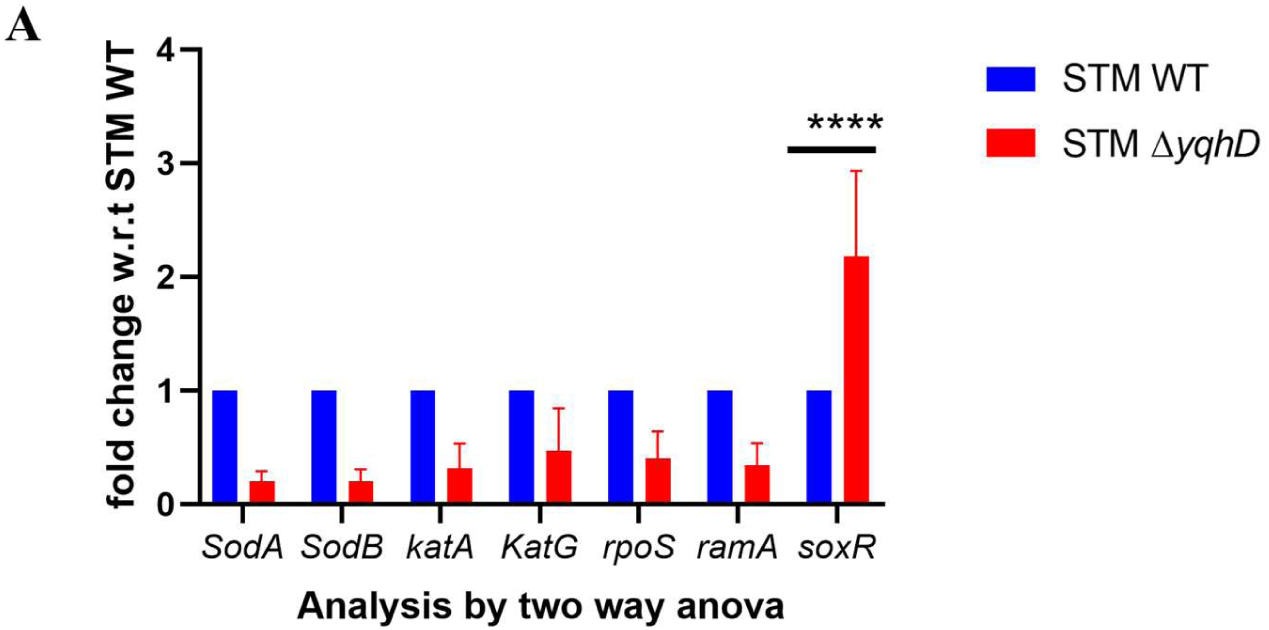
Antioxidant genes in STM Δ *yqhD* in HepG2 cells at 16 hours post-infection. Analysis was performed by using two -way ANOVA; p values****<0.0001, ***<0.001, **<0.01, *<0.05.

**Supplementary Figure 9.**
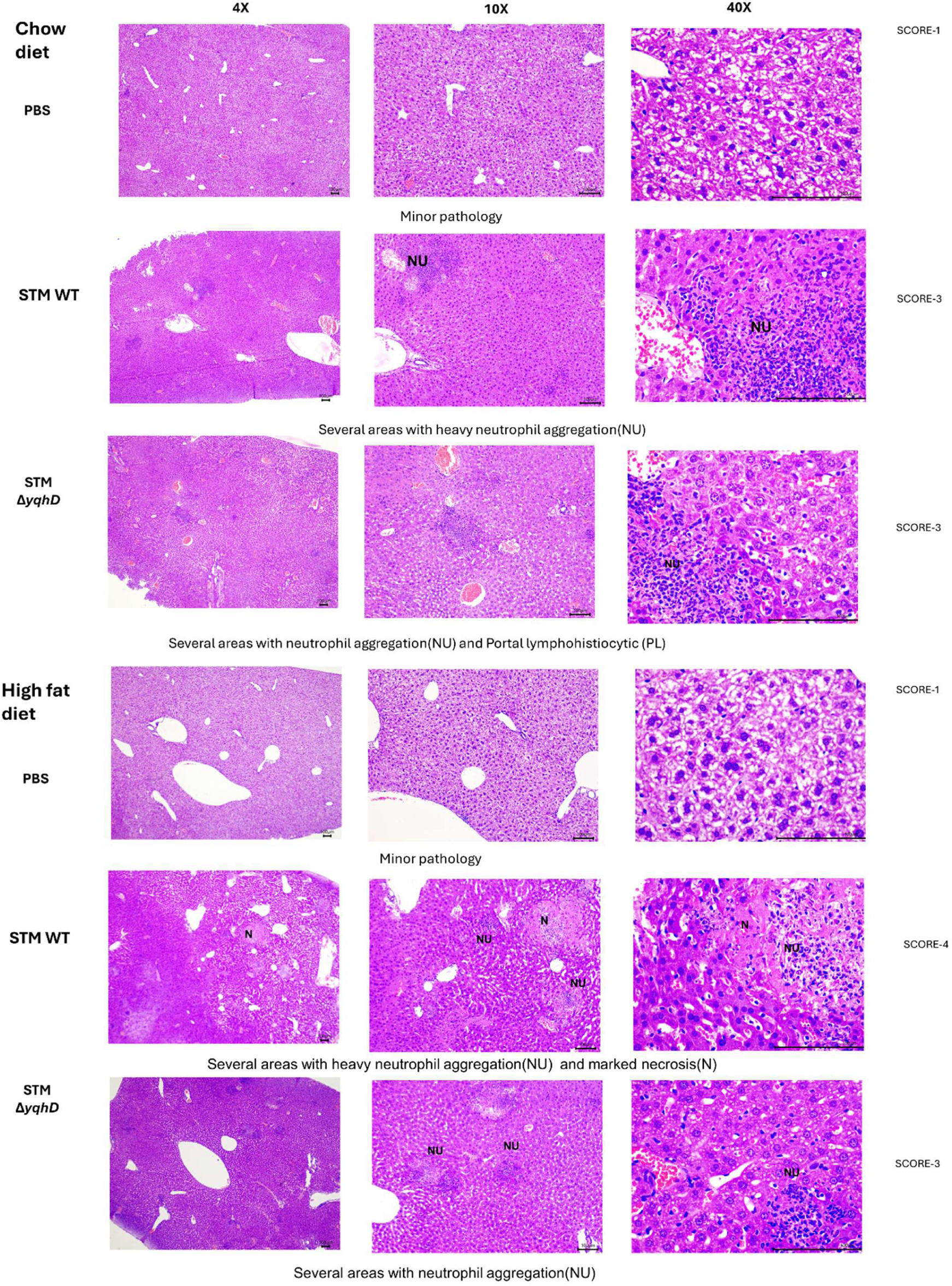
Haematoxylin and eosin staining of the liver sections from *gp91^phox−/−^* mice at various magnifications with the pathology score. Histopathology images of liver sections used in the main figure 4 at different magnifications with their pathology scores.

**Supplementary Fig 10.**
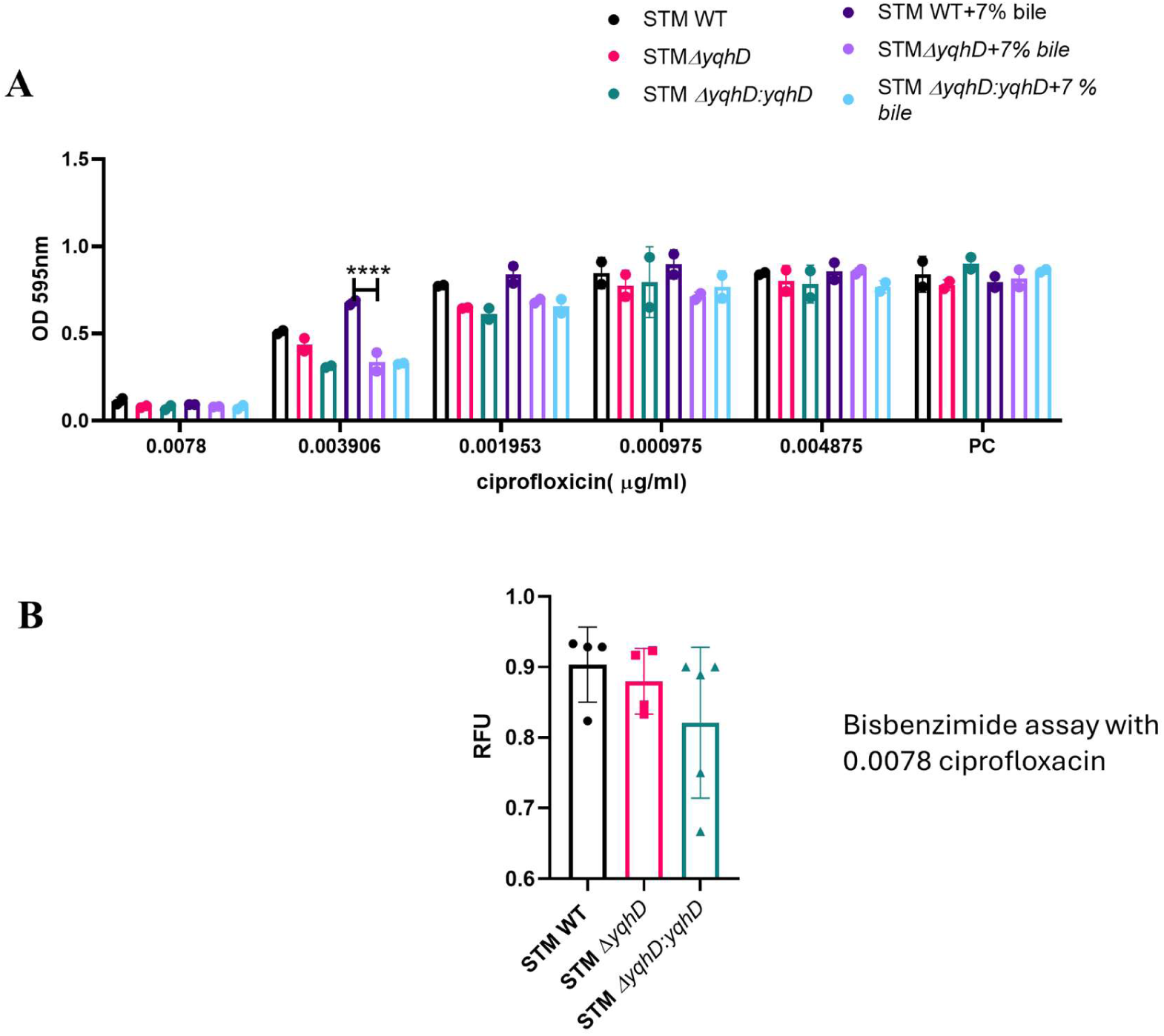
ciproflocin sensitivity increases on deletion of *yqhD* in *Salmonella*. **A.** Minimal inhibitory concentration (MIC) of STM WT, STM *ΔyqhD,* STM *ΔyqhD:yqhD* (N=3,n=2). Analysis was performed using two-way ANOVA. p values****<0.0001, ***<0.001, **<0.01, *<0.05. **B.** Bisbenzimide assay of STM WT, STM *ΔyqhD,* STM *ΔyqhD:yqhD* on treatment with 0.0078 µg/ml of ciprofloxacin(N=3,n=4)

**Supplementary Fig 11.**
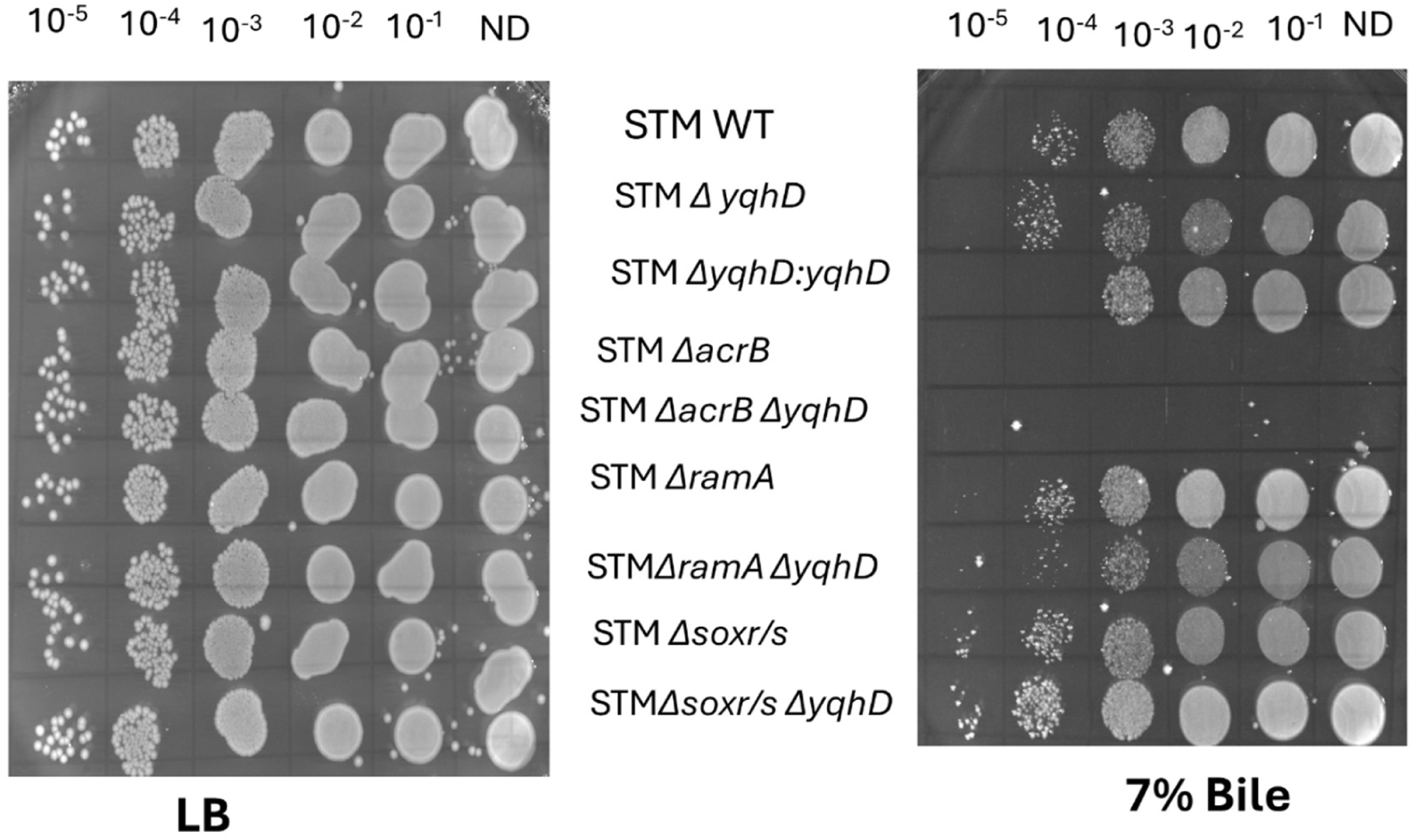
Spot assay on the plate with LB-agar or LB-agar supplemented with 7% bile. Spot assay of STM WT, STM *ΔyqhD,* STM *ΔyqhD:yqhD,* STM *ΔacrB,* STM *ΔacrB ΔyqhD,* STM *ΔramA,* STM *ΔramA ΔyqhD,* STM *ΔsoxR/S and* STM *ΔsoxR/S ΔyqhD* (N=3). Spot volume= 5 microlitres of 0.3 OD adjusted bacteria

**Supplementary Fig. 12.**
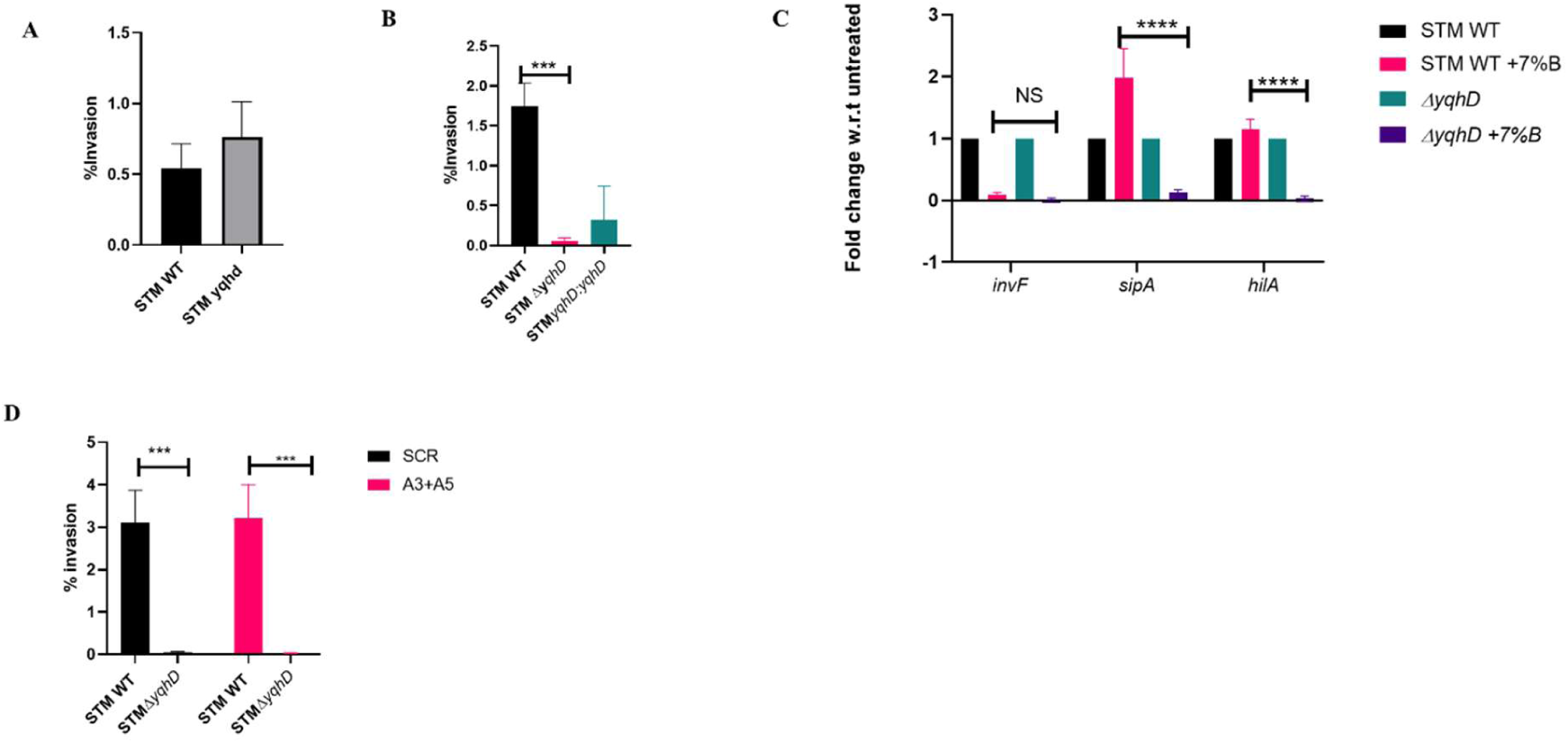
Deletion of *yqhD* in *Salmonella* decreases invasion. **A,B**. Invasion of STM WT, STM *ΔyqhD,* STM *ΔyqhD:yqhD* in colon carcinoma and HepG2 cells, respectively. Analysis performed using unpaired Student’s t-test. p values****<0.0001, ***<0.001, **<0.01, *<0.05. **C**. Q-RT PCR of invasion genes on bile salt treatment in STM WT, STM *ΔyqhD.* Fold change has been calculated with respect to untreated samples grown in LB. Analysis performed using two-way ANOVA.; p values****<0.0001, ***<0.001, **<0.01, *<0.05 **D**. Percentage invasion in HepG2 cells with CYP7A1 knockdown. Analysis performed using two-way ANOVA.; p values****<0.0001, ***<0.001, **<0.01, *<0.05

